# Synthetic gene networks recapitulate dynamic signal decoding and differential gene expression

**DOI:** 10.1101/2021.01.07.425755

**Authors:** Dirk Benzinger, Serguei Ovinnikov, Mustafa Khammash

## Abstract

Cells live in constantly changing environments and employ dynamic signaling pathways to transduce information about the signals they encounter. However, the mechanisms by which dynamic signals are decoded into appropriate gene expression patterns remain poorly understood. Here, we devise networked optogenetic pathways that achieve novel dynamic signal processing functions that recapitulate cellular information processing. Exploiting light-responsive transcriptional regulators with differing response kinetics, we build a falling-edge pulse-detector and show that this circuit can be employed to demultiplex dynamically encoded signals. We combine this demultiplexer with dCas9-based gene networks to construct pulsatile-signal filters and decoders. Applying information theory, we show that dynamic multiplexing significantly increases the information transmission capacity from signal to gene expression state. Finally, we use dynamic multiplexing for precise multidimensional regulation of a heterologous metabolic pathway. Our results elucidate design principles of dynamic information processing and provide original synthetic systems capable of decoding complex signals for biotechnological applications.

## Introduction

One of the most fundamental functions of living cells is the ability to respond to their environment by adjusting gene expression levels. Importantly, the signals that cells encounter may change on fast time-scales. These arise, for example, from passively changing stimuli, such as changes in nutrient concentration by diffusion, but also from actively coordinated signals in multicellular consortia and tissues. Examples for the latter are cyclic AMP oscillations during the aggregation of Dictyostelium [1] and the periodic/pulsatile secretion of hormones in mammals [2]. In addition to temporally changing inputs, an ever increasing number of studies show that signaling is highly dynamic, even if the received signals are constant [3].

How cells decode signal and signaling dynamics is not yet well understood [3]. One interesting hypothesis is that cells employ multiple downstream regulators that exhibit different response kinetics and thus respond selectively to certain dynamic input signals [4–9]. For example, it was shown that NFAT isoforms show distinct nuclear localization kinetics to cell stimulation and stimulus release [4,5]. Furthermore, decoding may occur on the level of promoter transitions of target genes [10,11] or on the level of gene regulatory networks (GRNs) [12]. Lastly, one of the most established ways cells may decode signals is by regulating multiple steps involved in target gene expression. For example, the immediate early gene c-fos is transcriptionally induced and stabilized in response to ERK activity which allows for the sensing of the duration of ERK activity [3,13]

The availability of synthetic systems that can process and distinguish fast dynamic signals would enable us to engineer cells that can precisely respond to their environment in defined ways and would shed new light on natural systems. Furthermore, multiplexing dynamic input signals would give us an increased capacity to instruct/communicate with cells for biotechnological and biomedical applications. To date, most synthetic biology circuits are based on gene regulatory interaction and have thus limited ability to react to information contained in fast dynamic signals. More recently, efforts were made to establish a toolbox for constructing synthetic signaling circuits in yeast [14]. These circuits were then combined with gene networks that act on slower time-scales, similar to natural cellular decision making circuits [14]. Furthermore, synthetic immediate early genes were constructed that can process dynamic ERK activity [13,15].

In this study, we set out to construct synthetic circuits in *S. cerevisiae* that can process and distinguish dynamic signals based on principles used in natural systems (**Fig. 1**). We decided to use optogenetic circuits as a prototype due to the ease of generating dynamic light signals. Specifically, we were interested in evaluating to which extent multiple regulators that act in parallel can be used to extract information about the strength and dynamics of input signals. For this, we constructed optogenetic transcriptional activators and repressors with different response kinetics and used them either as part of diamond-shaped incoherent feedforward loops (diamond-IFFLs) [16,17] or in isolation. We show that the diamond-IFFL acts as a falling edge detector and that its dose-response behavior to pulsatile and constant inputs can be employed to decode dynamically encoded signals—a property that we use to regulate the flux through a heterologous metabolic pathway. By combining the response of multiple target genes in GRNs, we show that systems can be created that allow for the distinction and decoding of constant (pulse-rejection filter) and intermittent (pulsatile signal decoder) signals. Our observations give insights into how cells decode dynamic signals and may guide the design of synthetic circuits that can process complex signals found in natural cellular environments.

**Figure 1.**
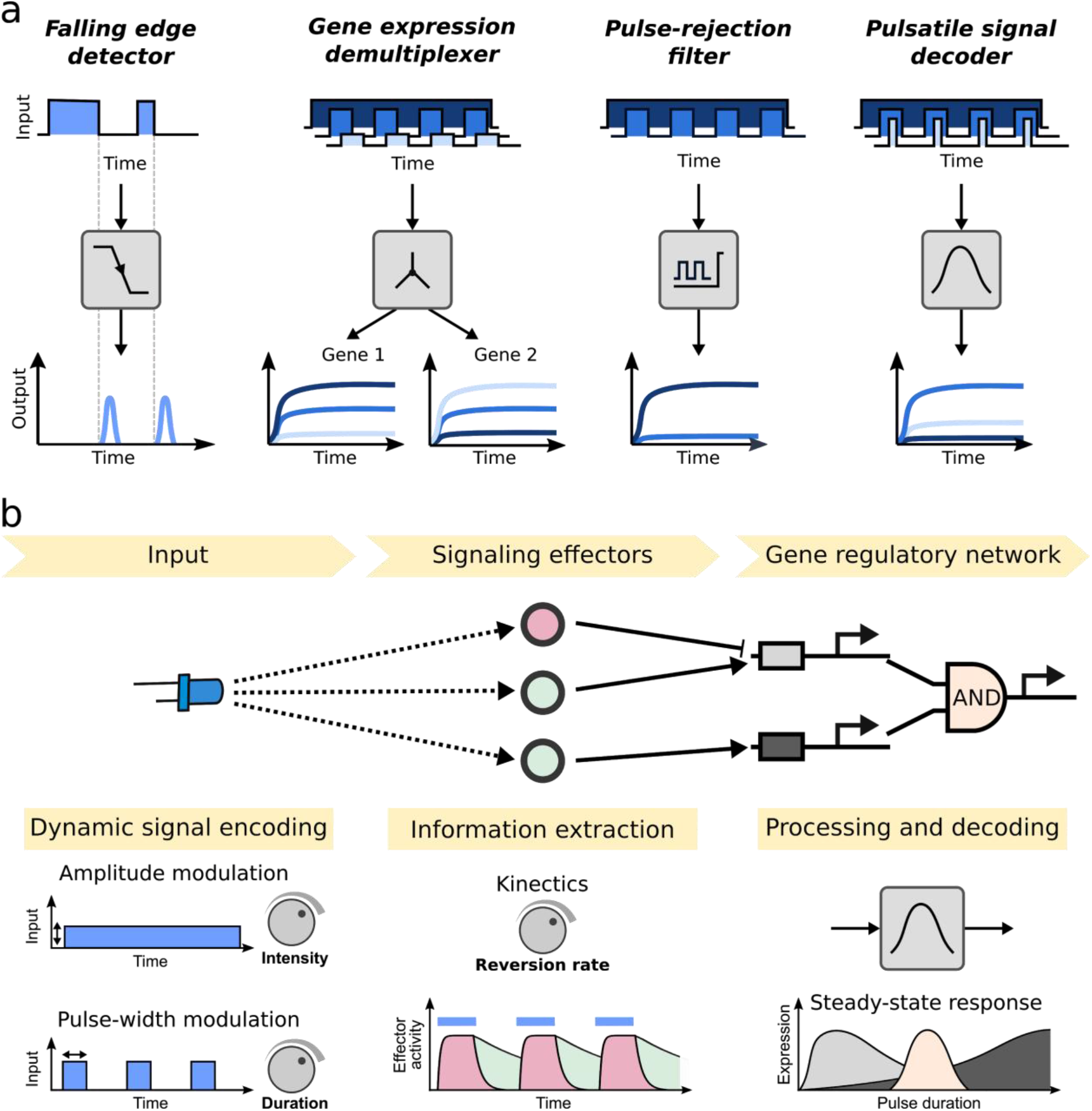
Synthetic circuits for dynamic signal processing and decoding. **(a)** Summary of dynamic signal processing functions. In this study, we construct a “falling edge detector” that, when triggered by the falling edge of an input pulse, generates a transcriptional pulse; a “gene expression demultiplexer” that enables differential expression of multiple genes based on a single dynamically multiplexed signal; a “pulse-rejection filter” that rejects intermittent signals while admitting continuous signals; and a “pulsatile signal decoder” that detects the presence of pulsatile signals and ‘reads out’ their pulse width duration. **(b)** Architectures for dynamic signal processing. Information can be encoded in signal and signaling dynamics. For example, in contrast to modulating the strength of a constant signal (amplitude modulation), the pulse duration of periodic pulsatile signals can be modulated (pulse-width modulation) (left). Upstream signal and signaling dynamics typically affect the activity of multiple signaling effectors, including transcriptional activators and repressors. These regulators may differ in their characteristics, such as the rate of reversion to an inactive state (middle). The activity of these regulators can in turn be integrated on the promoter level of direct target genes. The output of target genes can be further combined in gene regulatory networks which perform additional operations (right). This combined networked architecture can be configured to achieve signal processing functions, as shown in (a).

## Results

### Characterization of dark-state reversion mutants of EL222

To construct synthetic light-responsive circuits that can process dynamic signals, we first aimed to implement optogenetic transcription factors (TFs) with varying response kinetics. For many light-responsive protein domains, mutations are known that change the rate at which the protein switches from the photo converted to the dark state (dark-state reversion rate) [18]. One of these is the light-oxygen-voltage domain protein EL222 of *Erythrobacter litoralis* [19]. EL222 acts as a light-responsive DNA binding domain [19,20] and has been previously employed to regulate gene expression in mammalian cells, zebrafish, yeast, and bacteria [21–26]. Here, we constructed a set of TFs (referred to as Msn2AD-EL222) by fusing the Msn2 activation domain (AD) to the fast reverting wild-type EL222 (WT) and mutants with intermediate and slow dark state reversion rates (A79Q and AQTrip, respectively) [19] (**Fig. 2a,b**).

**Figure 2.**
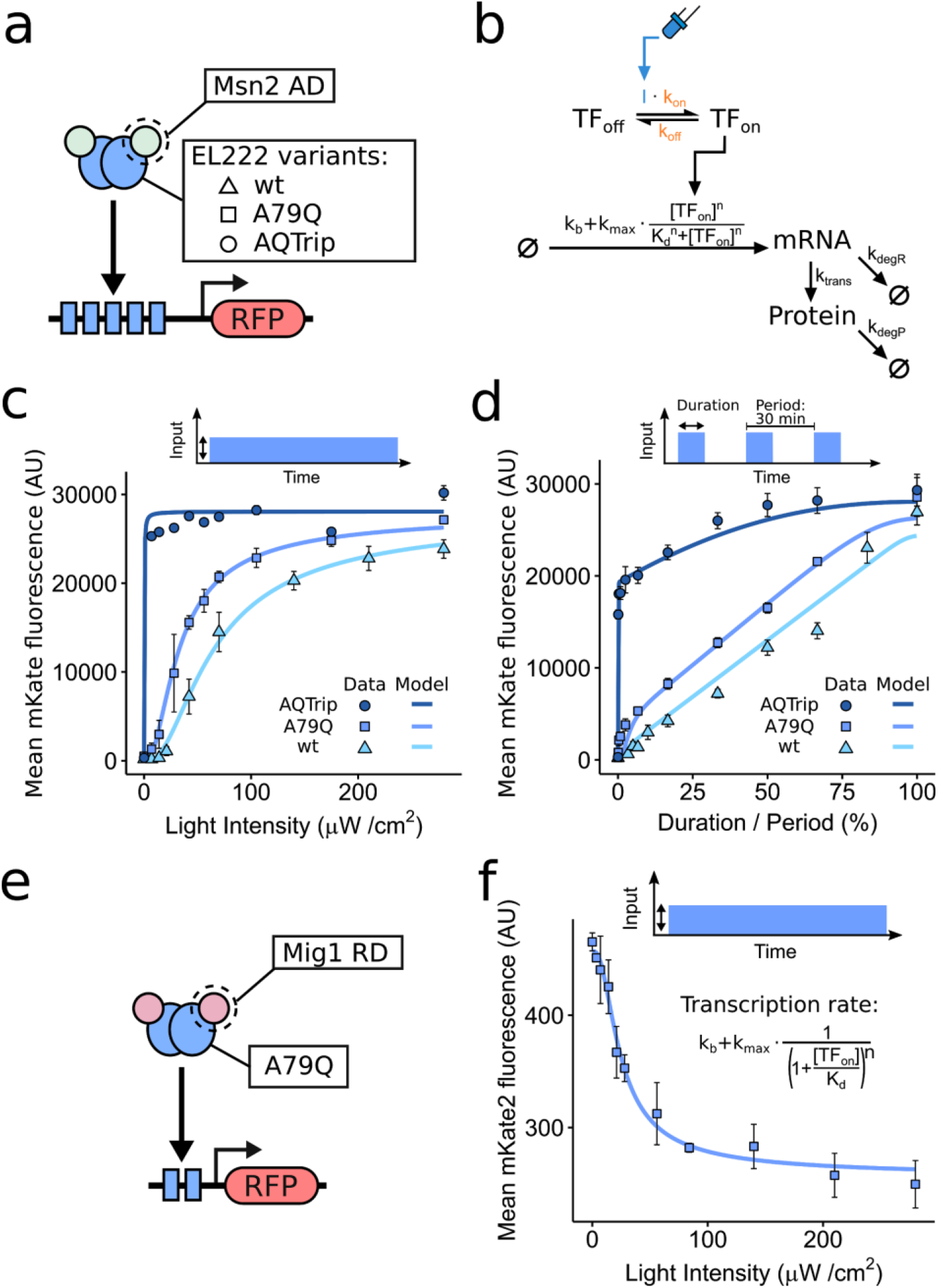
Characterization of optogenetic gene expression systems based on dark-state reversion mutants of EL222 in *S. cerevisiae*. **(a)** Schematic of EL222-based TFs and fluorescent reporter gene. The reporter gene consists of five EL222 binding sites followed by upstream of a truncated *CYC1* promoter and the coding sequence of the red fluorescent protein (RFP) mkate2 [25]. **(b)** Schematic of a model for Msn2AD-EL222 mediated gene expression [25]. ‘I’ represents the light input. k_on_ and k_off_ (orange) are allowed to vary between model descriptions of the dark-state reversion mutants. See Methods for details. **(c)** and **(d)** Dose-response of steady state reporter gene expression to amplitude modulation (AM) (c) and pulse-width modulation (PWM) (d) for different dark-state reversion mutants of EL222. Strains were grown under the illumination conditions depicted on the x-axis for 6 h. The light intensity and period for PWM were 280 µW cm^−2^ and 30 min. Data represents the mean and s.d. of two independent experiments. Lines represent model fits. **(e)** Schematic of the EL222-based transcriptional repressor and fluorescent target gene. The reporter gene consists of two EL222 binding sites followed by upstream of a truncated *CYC1* promoter and the coding sequence of the red fluorescent protein (RFP) mkate2 [25]. **(f)** Dose-response of steady state reporter gene expression in response to AM. The strain containing the constructs depicted in (a) was grown under the illumination conditions depicted on the x-axis for 6 h. Data represents the mean and s.d. of two independent experiments. Lines represent model fits. The model is that shown in Fig. 2b but with hill-type repression (equation shown in figure) instead of activation of transcription (Methods).

To evaluate the response characteristics of these TFs, we measured dose response curves of a mKate2 [27] fluorescent reporter gene [25] to different light intensities (referred to as AM for amplitude modulation) and durations of periodic signals (referred to as PWM for pulse width modulation) (**Fig. 2c-d**) [25]. We found that the AQTrip mutant is sensitive to low light intensities and is almost maximally induced by the lowest intensity tested. Both other mutants result in more sigmoidal dose-response curves with A79Q being more sensitive to light intensity than wt. As shown previously, the PWM modulation of the WT EL222-based TF (30 min pulse period) resulted in an almost linear dose response curve due to fast dark-state reversion [25]. In contrast, the responses of the dark-state reversion mutants are characterized by two phases - an initial steep response in gene expression to short light pulses, followed by a more gradual response that is close to linear for the A79Q mutant and slowly saturating for AQTrip.

To understand whether differences in the light response kinetics of the different TFs can indeed explain the observed behaviour on the gene expression level, we fit a simple mathematical model [25] describing TF activation and target gene expression to the data while only allowing conversion rates to vary between TFs (**Fig. 2b**, see Methods for details). We found that this model can recapitulate the data very well and recovered the expected ordering of photoactivated state half-lifes (wt ≈ 2 min < A79Q ≈ 6 min < AQTrip ≈ 27 min) (**Fig. 2c,d**). These rates stand in agreement with the lifetime of the photogenerated cysteinyl-flavin adduct in EL222 mutants measured in vitro (approximately 0.5, 5, and 33 min for wt, A79Q, and AQTrip) [19]. Thus, using dark-state reversion mutants of EL222, we were able to construct optogenetic TFs with different response characteristics.

### A diamond-IFFL acts as a falling edge detector

Having established optogenetic TFs with different response kinetics, we next aimed to use them to regulate target genes in a combinatorial manner. The most straight-forward design to accomplish this is a diamond-shaped incoherent feedforward loop [16] in which transcriptional activators and repressors act on the same target gene (**Fig. 1**). For this, we constructed an optogenetic transcriptional repressor [28] by fusing the Mig1 repression domain to EL222 (Mig1RD-EL222) (**Fig. 2e**). This domain was chosen because in *S. cerevisiae* Mig1 is known to co-regulate genes with the transcriptional activator Msn2 based on the relative timing of their activity, with target genes being repressed when both regulators are active [29]. We characterized the ability of this TF to repress leaky expression from EL222-responsive promoter based on a minimal *CYC1* promoter and two EL222 binding sites [25]. By measuring expression in response to AM, we found that the repressor is functional and inhibits gene expression in a dose-dependent manner (**Fig. 2f**).

Having established that the repressor is functional, we next integrated an activator with the AQTrip mutation and a repressor based on wt EL222 (the diamond-IFFL) into a single yeast strain and measured the transcriptional response of a target gene in live-cells by using the PP7 system (**Fig. 3a**) [30]. To increase the likelihood of dominant repression, Mig1RD-EL222 was expressed under the control of a stronger promoter (pr*TDH3*) [31] than Msn2AD-EL222 (pr*RPL18B*). For this we integrated a reporter gene consisting of five EL222-binding sites and a truncated *CYC1* promoter [25] as well as a sequence encoding 24 copies of the PP7 stem-loop upstream of the endogenous *GLT1* open reading frame [26]. A constitutively expressed fusion protein, consisting of a PP7 bacteriophage coat protein tandem dimer fused to an NLS and two copies of the red fluorescent protein mRuby3, binds to these stem-loops, allowing for the visualization of nascent RNAs as a fluorescent diffraction-limited spot in the nucleus [26]. We observed the cellular response to a single light pulse and found that there was only spurious transcription before and during the pulse (**Fig. 3b**). However, starting 4 min after the end of the light pulse, cells showed a strong transcriptional response that lasted for about 15 min. Importantly the post-pulse response is essentially identical between a 4 min and 30 min light pulse (**Fig. 3b**). These results show that the diamond-IFFL response is indicative of the end of a light pulse, similar to falling edge detectors used in signal processing and engineering [32].

**Figure 3.**
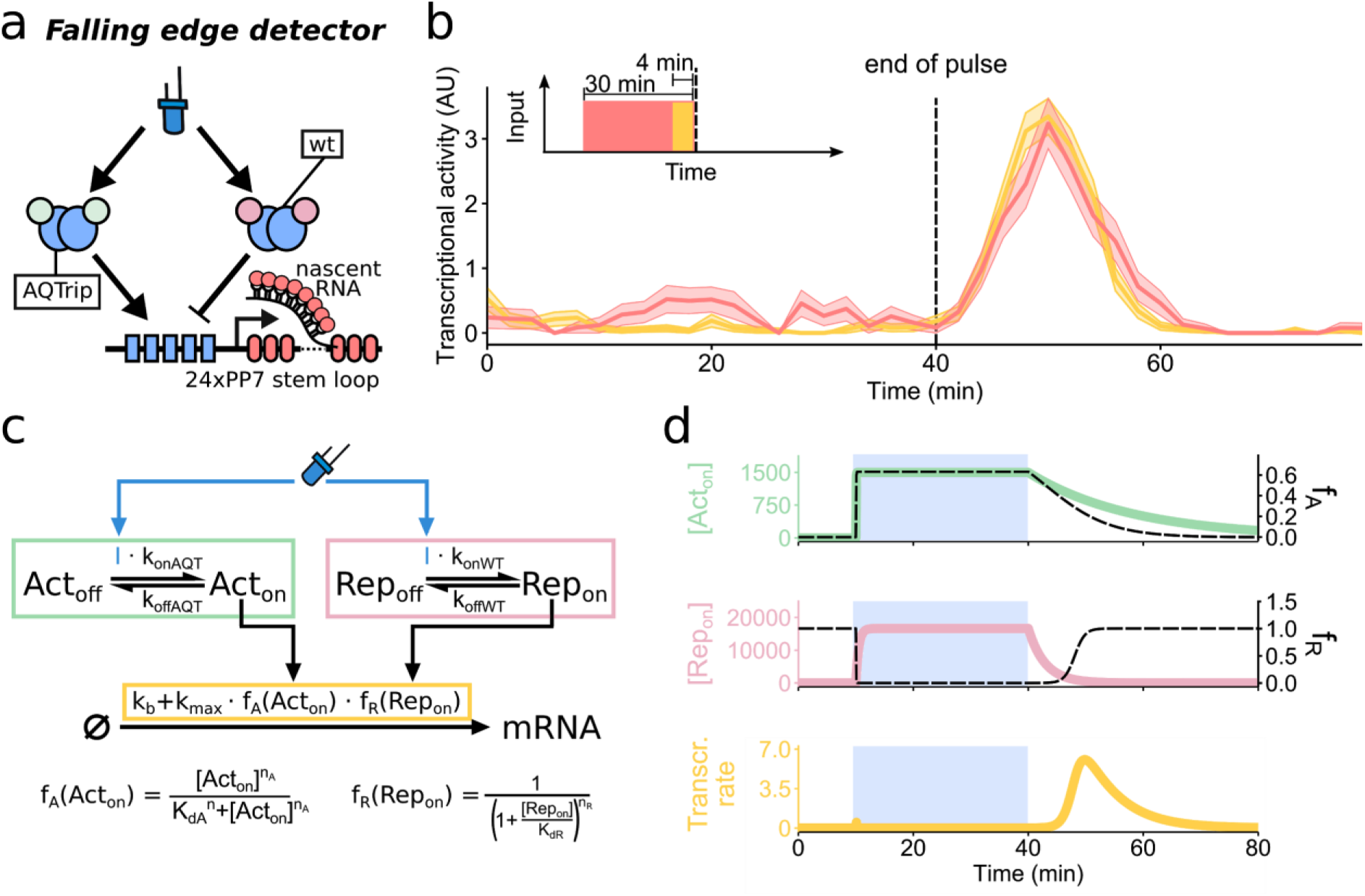
A diamond-IFFL acts as a falling edge detector. **(a)** Schematic of diamond-IFFL and the PP7-based transcriptional reporter. **(b)** Averaged single-cell transcriptional responses to a 4 min (yellow) or 30 min (red) blue light pulse. The light pulse ended at t = 40 min for both experiments and cells were not illuminated with blue light before and after the pulse. The line represents mean and the shaded area the s.e.m. for 171 to 250 cells. **(c)** Schematic of a model for diamond-IFFL-mediated gene expression regulation. The model describes Msn2AD-EL222 (green box) and Mig1-RD-EL222 (red box) regulation as well as their effect on transcription rate, modeled as the multiplication of an activating and repressive hill-function (yellow box). **(d)** Simulation results of the diamond-IFFL response to a 30 min light pulse (shown as blue background color). The temporal response of the activator (top, green), repressor (middle, red) and target gene transcription rate (bottom, yellow) are shown. Parameters are the ones derived from the characterization experiments in Fig. 2c,d, and f (see Methods for details). Only the total concentrations of the activator and repressor were calibrated to 1500 and 20000, respectively, to qualitatively recapitulate experimental results shown in (d). This adjustment is justified by the choice of promoters controlling activator and repressor expression (main text).

The circuit behavior can be explained based on the different reversion rates of the repressor and the activator and can be recapitulated by a simple model of diamond-IFFL mediated gene expression (**Fig. 3c,d**). Both TFs activate quickly at the onset of the pulse, resulting in a negligible transcriptional response due to dominant repression. After pulse termination, the repressor will return to its inactive dark-state within minutes while the activator will stay active for a longer time, resulting in a pulse of transcription whose length depends on the reversion rate of the repressor (**Fig. 3d**). One interesting property of such a falling edge detector is that the total transcriptional response should be independent of pulse duration as long as pulses have a distance from each other that is larger then the length of the transcriptional post-pulse response, meaning about 20 min for the diamond-IFFL [33]. Thus, this design may be used in the future to construct pulse counting circuits [33–35].

### The diamond-IFFL shows non-monotonic dose response curves

Having shown on the transcriptional level that the diamond-IFFL is functional and that the repressive function of Mig1RD-EL222 can be dominant over Msn2AD-EL222, we next characterized the steady state response to AM and PWM signals on the gene expression level. For this, we constructed four diamond-IFFLs with the AQTrip mutant activator and different repressor EL222 mutants and expression levels (**Fig. 4a**). We found that diamond-IFFLs show a non-monotonic dose-response to both types of input signals (**Fig. 4b,c**). Such behaviour was previously described for IFFLs in response to constant input signals [36,37]. The choice of EL222 mutant used in the repressor did not strongly affect the PWM response but resulted in a stronger gene expression response at intermediate light intensities in the case of regulation by AM. Similarly, increasing Mig1RD-EL222 expression levels resulted in a weaker AM response while not affecting the PWM response. Reducing the expression level of Mig1RD-EL222 resulted in failure to achieve the pre-induction state for maximal input signals. Furthermore, for PWM the response of this diamond-IFFL is maximal at intermediate pulse durations. These results show that by changing the kinetics and expression levels of its components, the response characteristic of the diamond-IFFL can be tuned. Importantly, using these modifications the response to constant (AM) and pulsatile (PWM) signals can be affected differentially.

**Figure 4.**
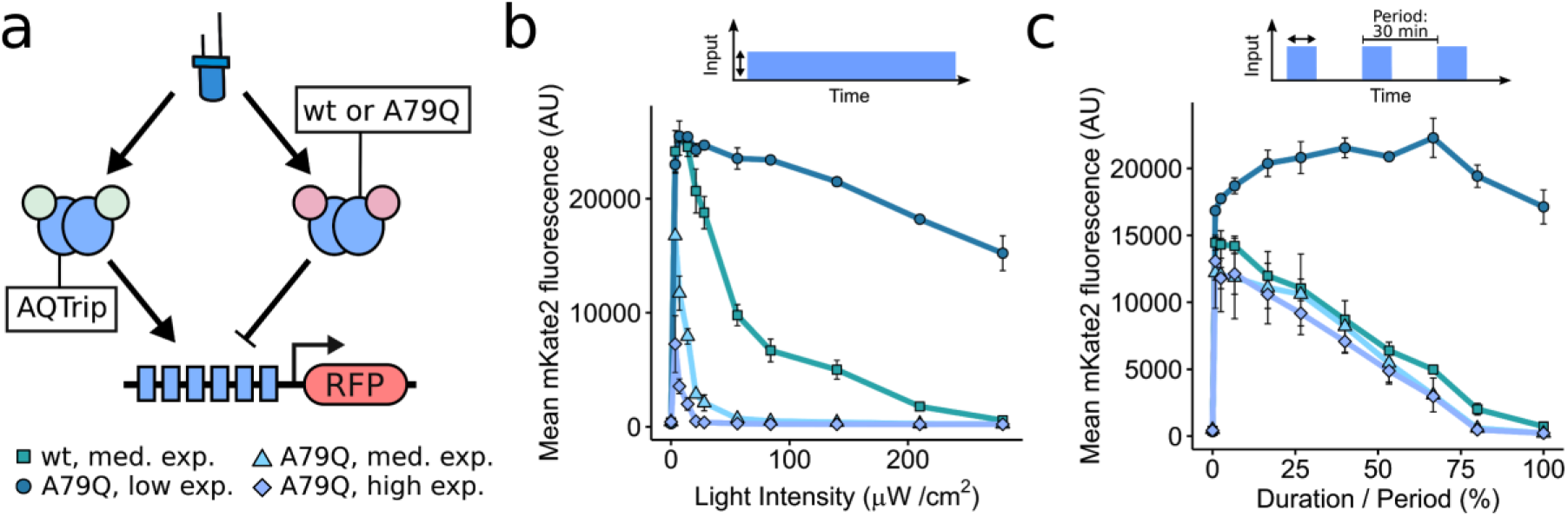
Diamond-IFFLs show tunable, non-monotonic dose-response curves. **(a)** Schematic of the diamond-IFFL and fluorescent target gene. IFFLs differing in repressor EL222 mutants and expression levels were constructed (the *TDH3* promoter was used for medium and the *HHF1* promoter was used for low expression [31], high expression was achieved using a second copy of the repressor gene under (*TDH3* promoter)). **(b)** and **(c)** Dose-response of steady state reporter gene expression to AM (b) and PWM (c) for diamond-IFFL variants (triangle = highly expressed wt-based repressor, square = highly expressed A79Q-based repressor, circle = medium expressed A79Q-based repressor). Strains were grown under the illumination conditions depicted on the x-axis for 6 h. The light intensity and period for PWM were 280 µW cm^−2^ and 30 min. Data represents the mean and s.d. of two independent experiments.

### Combination of optogenetic systems allows for dynamic multiplexing

In biological systems, dynamic encoding is thought to allow for the transmission of complex information, such as the identity and strength of a stressor, in a single time-varying signal [3,38–41]. The decoding then allows to differentially regulate genes based on the information contained in the signal [3,10,40,42]. Having synthetic systems that enable such regulation would allow us to easily control the expression of various genes and cellular processes independently. This may be especially important for optogenetic regulation, as overlap in absorption spectra of light-responsive as well as fluorescent reporter proteins limits the use of multiple regulators in a single cell without crosstalk [43,44].

We thus tested whether we could achieve multiplexing with a single dynamic light signal by combining the non-monotonic diamond-IFFL with a second blue-light responsive gene expression system. For this, we used a TF (referred to as liLexA hereafter) based on the light-regulated heterodimerization of *Arabidopsis thaliana* cryptochrome 2 (CRY2) fused to the DNA binding domain and its binding partner CIB1 fused to the VP16 activation domain [45–47]. To enable a graded response to PWM [25], we used a fast dark-state reversion mutant of CRY2 (W349R) [45].

We added the liLexA construct together with a yellow fluorescent protein (YFP) reporter gene into a strain that contained the diamond-IFFL (based on a highly expressed A79Q-based repressor, see **Fig. 4**) with a RFP reporter gene (**Fig. 5a**). Measuring YFP expression in response to AM signals, we found that the liLexA system shows a sigmoidal dose-response curve that has only little overlap with RFP expression mediated by the diamond-IFFL (**Fig. 5b**). In contrast, PWM regulation of liLexA resulted in a close to linear dose-response curve due to the fast dark-state reversion of the W349R mutant [25,45] (**Fig. 5c**). As a consequence of this and the non-monotonic dose-response curve of the diamond-IFFL, YFP and RFP expression coincides at intermediate pulse durations and the expression ratio of both proteins changes with pulse duration. Lastly, we performed PWM with low light intensities and found that under these conditions, RFP is expressed in a graded fashion while YFP expression stayed approximately constant (**Fig. 5d**).

**Figure 5.**
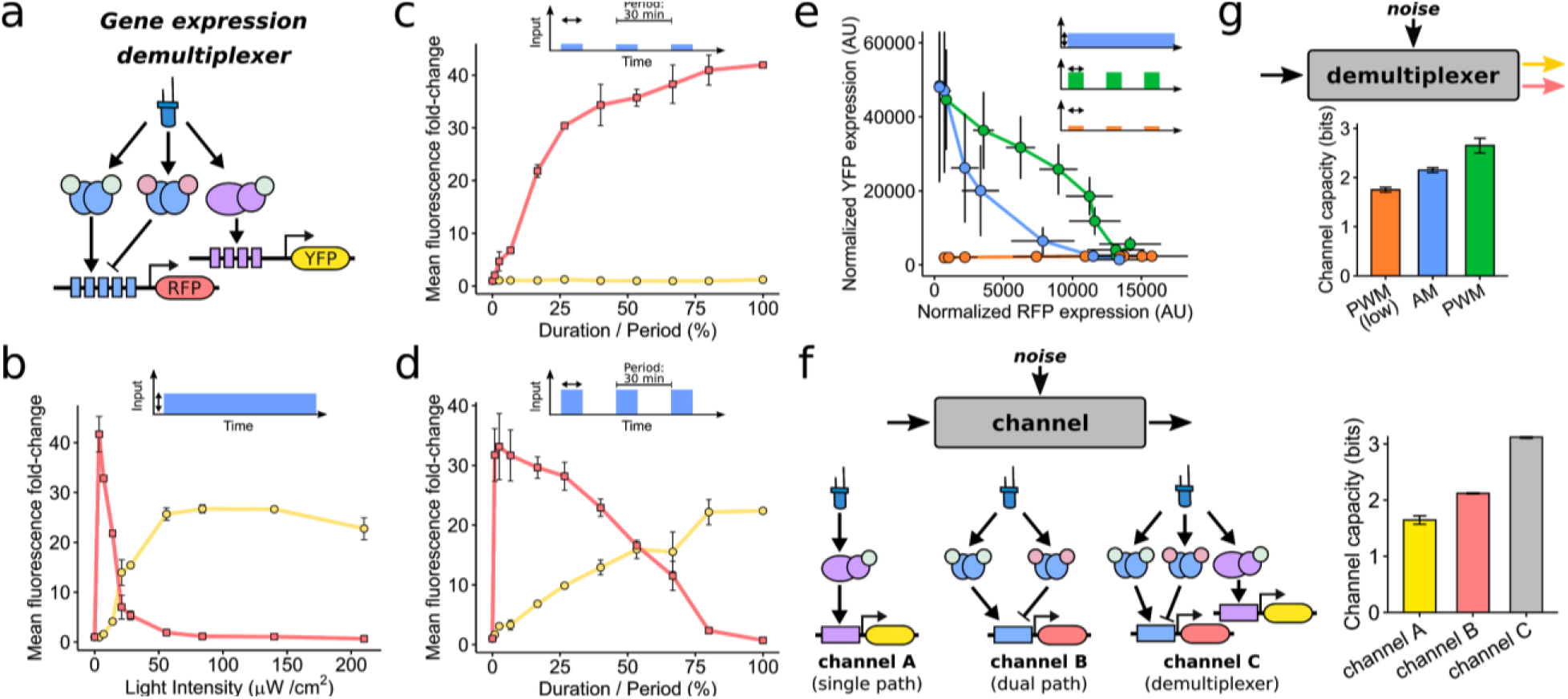
Independent regulation of two genes using a single, dynamically encoded signal. **(a)** Schematic of dynamic multiplexing circuit based on a diamond-IFFL and a liLexA-based expression system. **(b-d)** Dose-response of steady state YFP (circles) and RFP (squares) reporter gene expression to AM (b), low intensity PWM (c), and high intensity PWM (d). Strains were grown under the illumination conditions depicted on the x-axis for 6 h. Fluorescence fold-changes compared to dark conditions are shown. The light intensity for high intensity PWM was 210 µW cm^−2^ (c) and 3.5 µW cm^−2^ for low intensity PWM (d). The PWM period was 30 min. Data represents the mean and s.d. of two independent experiments. (**e**) Two-dimensional gene expression responses for different input signal types (blue = AM, green = high intensity PWM, orange = low intensity PWM). Data was taken from experiments shown in (c)-(d) and represents the mean and s.d. of measured fluorescent distributions normalized by side-scatter measurements. Responses for no and maximal input signals are not shown. (**f-g**) Information theoretic analysis of the gene expression demultiplexer. Channel capacity was estimated for three different channels, namely the demultiplexer and its individual branches (f, left), using all three input types (f). Estimation results of demultiplexer channel capacity using either of the three input types are shown in (g). Results represent the mean and s.d. calculated based on two independent experiments. See Methods for details on channel capacity calculation.

### Information theoretic analysis of the gene expression demultiplexer

By evaluating the two-dimensional gene expression responses of the demultiplexer, we found that the three input types (AM and low/high intensity PWM) enable access to different areas of the response space (**Fig. 5e**). Given that noise in signal sensing/transduction and gene expression can deteriorate cellular signal discrimination [48–50], we aimed to evaluate how accurately information is transmitted by the demultiplexer. Recently, information theoretic analyses were used to quantify the reliability of information transmission in cellular systems by treating signaling pathways or promoters as communication channels and calculating the maximal mutual information (referred to as channel capacity C) between input and output [41,48,49,51] (Methods). The value 2^C^ can be interpreted as an upper bound on the number of distinct input signals that can be transmitted by the channel without error.

Using this approach and a previously published algorithm [52], we quantified information transmission by the demultiplexer (**Fig. 5f, g**). Taking the response to all three input types into account, we found that the individual branches of the demultiplexer, meaning direct gene regulation by liLexA and dual path regulation by the diamond-IFFL, transmit limited information about the input signal with a channel capacity of 1.65 ± 0.08 (bits) and 2.15 ± 0.05 (bits), respectively, consistent with previous results [49] (**Fig. 5f**). However, the combined two-dimensional output of the demultiplexer results in a significant increase of channel capacity to 3.12 ± 0.02 bits (**Fig. 5f**, see also **Supplementary Fig. 1**) [48]. By restricting the types of inputs used, we further found that transmission reliability depends on the input type, with high intensity PWM resulting in the highest channel capacity (2.65 ± 0.15 bits) (**Fig. 5g**). This result is in agreement with our previous observation that PWM results in less cell-to-cell variability in gene expression compared to AM [25]. Importantly, jointly using all three input types, results in a higher channel capacity than using input types individually (**Fig. 5f,g**). Together with the finding that signaling dynamics carry more information of upstream signals than static responses [41,51], our data strongly indicates that dynamically multiplexed signals can be beneficial for accurate information transmission.

These results show that by employing a demultiplexer, two genes can be differentially regulated based on the strength and dynamics of a single input signal and that the use of such dynamic signals enables reliable information transmission.

### GRNs allows for further processing of the dynamic signals

Above, we described that dynamic multiplexing allows for the differential regulation of two target genes. The relative expression levels of those genes contain information about the strength and dynamics of input signals. However, to make this information generally accessible to cellular processes, the output of both target genes needs to be combined. For example, one may want to construct systems in which a gene of interest preferentially responds to sustained or intermittent signals. In many natural systems, direct target genes of a signaling pathway are further connected in gene regulatory networks [53]. Among others, the regulations in GRNs contain repression between target genes and combinatorial regulation of downstream genes [53]. To analyze whether we can use such networks to further process the outputs resulting from dynamic signals, we constructed synthetic gene regulatory interaction using dCas9-based transcriptional repressors and activators [54].

First, we sought to build a pulse-rejection filter by employing a network in which the diamond-IFFL output inhibits the expression of the liLexA target gene. We achieved this by expressing a synthetic guide RNA (sgRNA), which targets the LexA binding site, under control of the diamond-IFFL while expressing the repressor dCas9-Mxi1 [54] constitutively (**Fig. 6a**). By regulating the resulting strain using PWM, we found that at steady state the system only responds to long pulse durations (**Fig. 6a**). Thus the additional regulatory interaction allows for the distinction between constant (and close to constant) input signals and intermittent input signals.

**Figure 6.**
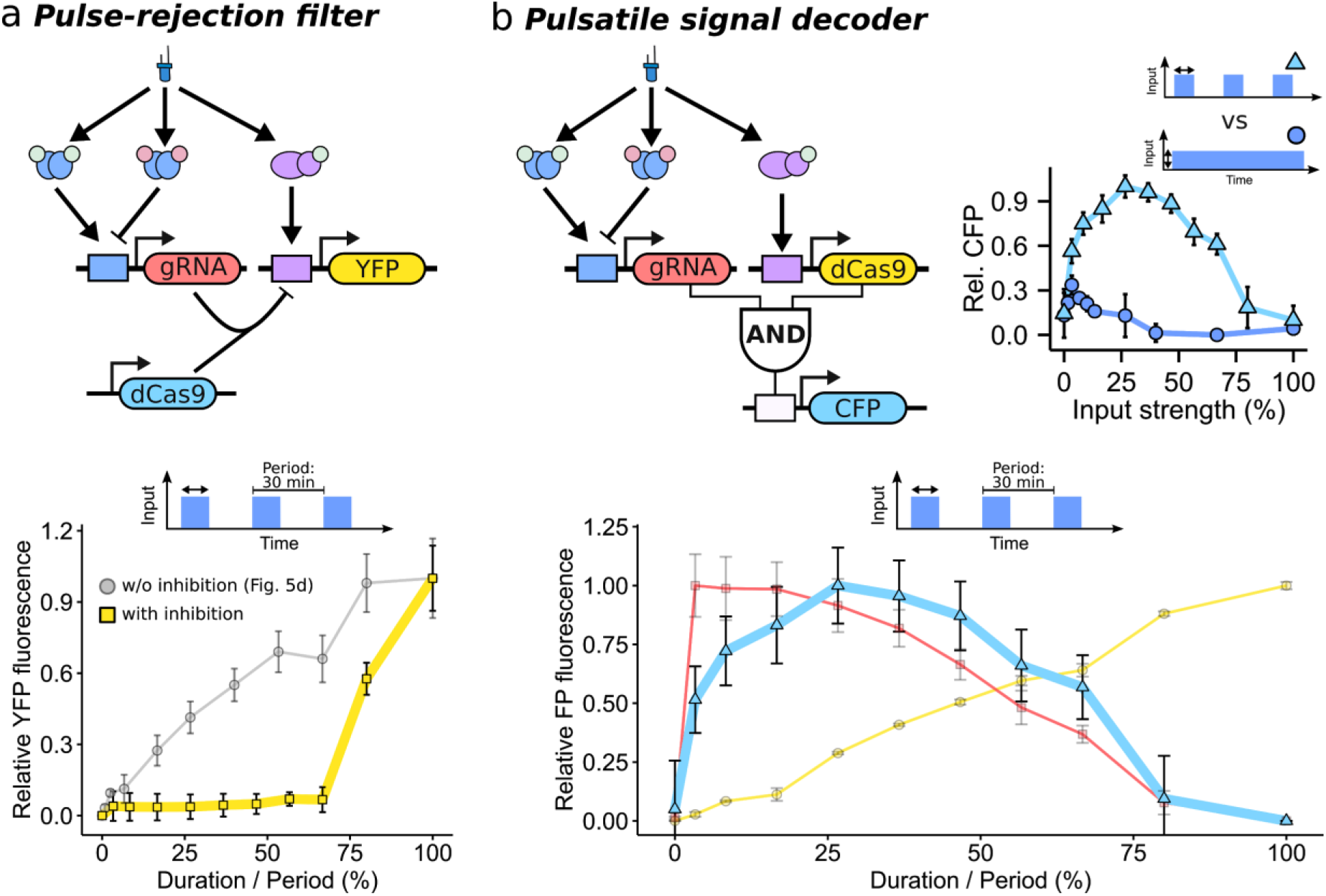
Circuits for detecting pulsatile inputs and decoding their pulse duration. **(a)** Pulse-rejection filter based on dCas9-Mxi-mediated repression of the liLexA target gene by the diamond-IFFL. A schematic of the circuit is shown on the top. Steady state dose-response of YFP expression in response to PWM of a strain containing the pulse-rejection filter (squares) and a control strain that only contains the diamond IFFL (circles, data is taken from Fig. 5d) is shown on the right. Cells were grown under the illumination conditions depicted on the x-axis for 12 h (repression system) or 6 h (control strain). Fluorescence values were normalized to lie between 0 (background expression) and 1 (maximal expression). The light intensity and period for PWM were 210 µW cm^−2^ and 30 min. **(b)** Pulsatile signal decoder based on a logical AND-gate combining the activity of the diamond-IFFL and liLexA (schematic, top left). Steady state dose response of the AND-gate strain to PWM (CFP, cyan triangles) is shown on the bottom. In addition to the constructs depicted in (c), the strain contains reporters for diamond-IFFL-mediated (RFP, red squares) and liLexA-mediated (YFP, yellow circles) gene expression. Top right shows a comparison of the steady-state response of the AND-gate strain in response to AM (blue, circles) and PWM (red, squares). Fluorescence levels were normalized to the minimal and maximal expression achieved between both types of regulation. Data of all experiments represents the mean and s.d. of two independent experiments.

Next we investigated the processing capabilities by combinatorial AND-gate regulation downstream of the diamond-IFFL and liLexA systems. For this, we put a sgRNA targeting the *GAL7* promoter under diamond-IFFL and the transcriptional activator dCas9-VP64 [54] under liLexA regulation (**Fig. 6b**). Furthermore, we added a reporter gene consisting of the *GAL7* promoter followed by the coding region of the cyan fluorescent protein (CFP) mTurquoise (**Fig. 6b**). Regulating this system with PWM, we found a non-monotonic dose-response curve of CFP expression which is maximal at intermediate pulse durations, where there is an overlap in the activity of diamond-IFFL and liLexA (**Fig. 6b**). Given that the overlap of both systems differs between input signal types (**Fig. 5**), we next asked how this affects the output of the AND-gate to AM. We found that AM results in only moderate CFP induction that is lower than that achieved by PWM at almost all input levels (**Fig. 6b**). Thus, using AND-logic, we built a pulsatile signal decoder that detects the presence and duration of pulsatile signals.

These experiments show that simple dCas9-based GRNs allow for the extraction of information contained in direct target genes of optogenetic systems, thus enabling signal dynamics to be decoded in the level of a single output gene.

### Regulation of metabolism by multiplexed optogenetic systems

Bioproduction is among the most promising avenues for the technological application of synthetic biological circuits [55]. To optimize metabolite production, it is typically necessary to adjust the expression levels of many genes to appropriately redirect metabolic flux [56–58]. Furthermore, the need for dynamic pathway regulation is increasingly recognized, for example to balance the trade-off between growth and production during a biotechnological process [24,59–61]. Thus, having robust tools that enable the precise and timed regulation of various genes would be of great use for metabolic engineering. Optogenetics is ideally suited for this purpose and has been recently applied to improve isobutanol production by pulsed regulation [24]. However, due to spectral overlap optogenetic tools cannot be used for metabolic control to their full potential.

We thus aimed to test whether dynamic multiplexing can be used to differentially regulate the expression of metabolic enzymes. For this, we chose to focus on a heterologous beta-carotene pathway in yeast (**Fig. 7a**) [61,62]. Previous studies in yeast have found that expression levels of pathway enzymes affect the color of yeast cultures which represents differences in metabolic pathway flux [57,62]. We constructed a strain in which expression of the enzyme crtI is regulated by the diamond-IFFL, while the expression of crtYB and crtE (all from *Xanthophyllomyces dendrorhous*) is under liLexA control (**Fig. 7b**). Additionally, the catalytic domain of the *S*.*cerevisiae* HMG-CoA reductase HMG1 (tHMG1) was constitutively overexpressed to increase the flux to the pathway [62].

**Figure 7.**
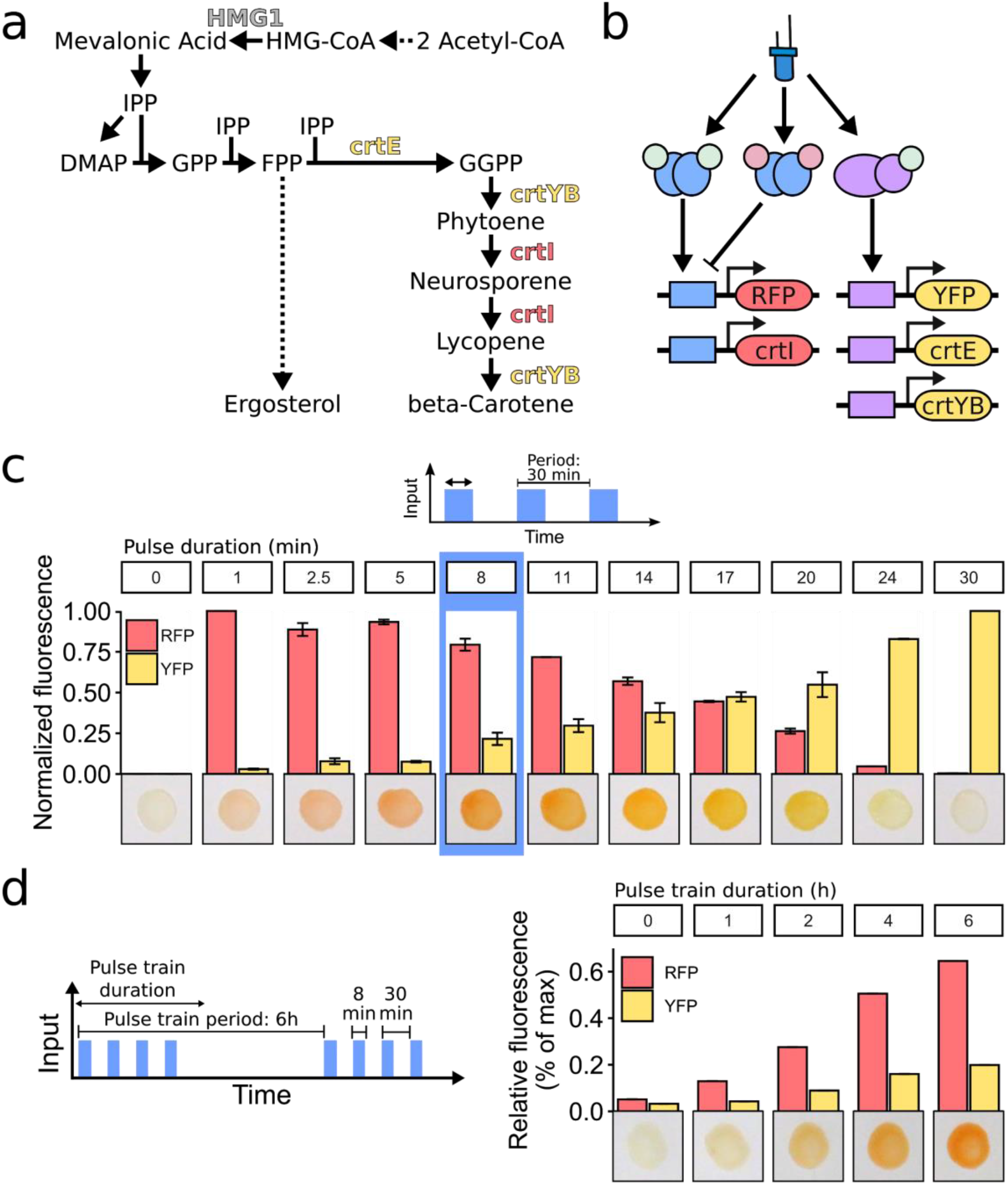
Regulation of a metabolic pathway with dynamic multiplexed signals. **(a)** Schematic of the heterologous beta-carotene pathway in *S. cerevisiae*. **(b)** Schematic of dynamic multiplexing system and fluorescent as well as metabolic target genes. **(c)** Fluorescent and metabolic response of a strain containing the system shown in (b) to PWM based regulation after 12 h of growth. Bars represent fluorescence level normalized to basal and maximal expression. Error bars represent the s.d. of two independent experiments. Images depict *S. cerevisiae* liquid cultures spotted and dried on filter paper (see Methods). PWM was performed with 210 µW cm^−2^ blue light intensity and a 30 min period. (**d**) Fluorescent and metabolic response to the modulation of pulse train duration. During the pulse train, pulse period was 30 min, pulse duration was 8 min, and blue light intensity was 210 µW cm^−2^. Fluorescence values are relative to the maximal value for each fluorescent protein measured in the experiment shown in (c).

When performing expression modulation of the resulting strain using PWM, we found that the different levels and ratios of enzyme expression resulted in color production ranging from faint rose to orange and yellow and then to faint yellow (**Fig. 7c**). In the absence of light as well as in the presence of constant light, yeast cultures appeared white. This data indicates that the regulatory overlap of the optogenetic systems results in an input dependent spectrum of products. Furthermore, the pathway can be seen as an AND-gate architecture at the metabolic level, in which carotenoid production only occurs when both gene expression systems are active.

The use of light as an input allows not only for the generation of PWM and AM signals, but also for the generation of dynamics of essentially arbitrary complexity. We hypothesized that periods of transcriptional inactivity can be used to tune the level of metabolite production. Thus, we decided to test whether the metabolic pathway flux can be tuned based on regulating the duration of periodic pulse trains—dynamics similar to periodic bursts of spiking in neurons [16,63] (**Fig. 7d left**). As hypothesized, by performing such a regulation, we found a modulation of color intensity rather than identity (**Fig. 7d**). The yeast color (**Fig. 7d,e**) is in accordance with the pure PWM experiments (compare **Fig. 7c**). Importantly, modulation of pulse train duration resulted in hues not achieved by PWM.

These results show that dynamic multiplexing can be used to perform multidimensional metabolic regulation based on a single input signal.

## Discussion

In this study we presented synthetic optogenetic circuits that allow for the processing and decoding of dynamic input signals based on multiple transcriptional effectors acting in parallel and gene regulatory interactions.

For this, we generated and characterized optogenetic transcriptional activators and repressors based on EL222 dark-state reversion mutants (**Fig. 2**). These TFs themselves represent valuable tools for gene expression regulation in basic research and synthetic biology applications. For example, TFs based on the AQTrip mutant are sensitive to low light intensities and may be used to avoid phototoxicity in scenarios where minute-scale regulation of transcription is not required, such as bioprocessing.

By following transcriptional output in live cells over time, we showed that a diamond-IFFL based on optogenetic TFs can act as a falling edge detector when the dark-state reversion rate of the activator is slower than that of the repressor (**Fig. 3**). A similar design based on other types of effectors can likely be achieved. For example, one can imagine tuning deactivation rates in phosphorylation based circuits [7,14], making the dynamic activity of many signaling pathways accessible to synthetic dynamic signal processing. The output of the falling edge detector can in the future be used to drive the expression of proteins that enable pulse counting and recording, such as recombinases and genome editors [34,35,64]. We show here that the same signal can act as the input to both the diamond-IFFL and a TF that directly affects expression of a second gene by itself (**Fig. 5**). Recording the output of multiple such target genes could give us detailed insights about the dynamic signals that cells experience in complex environments.

In nature, a form of pulse counting was proposed for the transcriptional response to cAMP waves in *Dictyostelium discoideum* [65]. There, cAMP results in the activation and - with a delay - in the nuclear export of the GATA TF GtaC [65]. This incoherent regulation of TF activity results in a response to the rising edge of the cAMP pulse. It will be interesting to investigate whether falling edge detectors based on our synthetic design can be found in nature. The exact implementation of the diamond-IFFL in this study is inspired by the coordinated gene regulation by the stress-responsive TFs Msn2 and Mig1 in *S. cerevisiae* [29]. The regulation of these TFs is highly dynamic, but to date gene expression responses were mainly investigated to steps in input signals, and it will be interesting to see how the relative activity of both TFs affects gene expression in response to more dynamic signals [29].

The dynamical behaviour of many signaling pathways is thought to encode information about the strength, dynamics, and identity of the signals they respond to [3,66]. However, for most pathways, the mechanisms which cells use to decode such signals and thus get access to this information are still not well understood [3]. Work on calcium and insulin signaling suggests that decoding may be achieved by employing signaling pathway branches with different response kinetics [4–6,8]. By building synthetic systems, we were able to show that signal decoding can be achieved by tuning the response characteristics of transcriptional effectors and combining their activity through gene regulatory interaction (**Fig. 5 and 6**). Different signaling proteins that exhibit dynamic activity, such as ERK [67], affect both the activity of transcriptional activators and repressor, and it would be interesting to investigate whether the response characteristics of those target effectors enable signal decoding similar to what we present here.

Studies using information theoretic approaches to investigate signaling fidelity found that dynamic signaling responses carry significantly more information than static ones [41,48] and that the dynamic response of multiple TFs can accurately represent complex extracellular environments [51]. However, there is still a limited understanding of whether (and how) dynamic signals can be interpreted reliably on the gene expression level [49]. For the yeast TF Msn2, it was found that constant TF activity is more reliably transmitted to target gene expression than pulsatile activity and that even the combination of multiple input types and target genes results in a maximal channel capacity of 1.8 bits [49]. The latter result was interpreted as Msn2 only allowing for the error-free transmission of signal type but not intensity [49]. However, the authors also showed that mutant promoters respond with higher fidelity than natural yeast promoters, suggesting that limited information transmission is not fundamentally given by the underlying molecular processes [49]. Using a synthetic gene expression demultiplexer, we show that the use of dynamic signals and multiple regulators allows for a significantly higher information transmission capacity from input signal to gene expression output of about 3.1 bits (**Fig. 5f**). We further found that pulse-width modulated signals can be transmitted more reliably than constant signals (**Fig. 5f**), showing that which signal type is optimally transmitted depends on the system in question [49]. Importantly, the synthetic circuit was not purposefully optimized for low gene expression variability, suggesting that our findings do not represent an upper limit of decoding fidelity.

The decoding ability allows us to send single multiplexed light input signals to control gene expression of multiple target genes (**Fig. 5**). Multiplexing is especially interesting for light-based regulation of cellular function, as the spectral overlap of different light-responsive proteins as well as fluorescent proteins may prohibit the independent modulation of distinct processes [44]. Furthermore, optogenetic regulation is not straightforward in organisms whose normal physiology requires ambient light, such as plants [43]. Dynamic signal encoding may prove useful in such scenarios, and we anticipate that our decoding circuits can be implemented in various organisms given its simple design based on widely applied light-responsive proteins [21–26,45,47]. Chemical production by metabolic engineering represents an application where the dynamic and independent regulation of many individual genes may open new avenues [57,59,60]. We show here that dynamic signal encoding can be employed to regulate the flux through a metabolic pathway (**Fig. 7**).

Taken together, our work shows how TFs with different response characteristics and simple gene regulatory networks can be used to multiplex, process, and decode dynamic signals and shows how these properties may be employed in a biotechnological context. These results give insight into how signal multiplexing and decoding may be achieved in natural systems, and will guide the design of synthetic circuits that can process complex signals found in cellular environments.

## Acknowledgments

We thank Victor Serban for technical assistance at an early stage of the project and members of the Khammash lab for helpful discussions. We would further like to thank Fabian Rudolf (ETH Zurich), Chandra Tucker (UC Denver), John Dueber (UC Berkeley), Stanley Qi (Stanford University), and Junbiao Dai (Shenzhen Institute of Synthetic Biology) for providing plasmids.

## AUTHOR CONTRIBUTIONS

D.B. conceived the study, performed experiments and mathematical modeling, and analyzed data together with M.K. S.O. performed experiments and mathematical modeling. M.K. supervised the study and secured funding. D.B. wrote the manuscript with contributions from M.K.

## COMPETING INTERESTS

The authors declare no competing interests.

## Methods

### Plasmid construction

*E. coli* TOP10 cells (Invitrogen) were used for plasmid cloning and propagation. All plasmids were constructed using the MoClo yeast toolkit standard using the procedure described in [31]. All parts used to construct final plasmids are summarized in **Supplementary Table 1** and all plasmids used in this study are listed in **Supplementary Table 2**. All part plasmids were verified by sanger sequencing (Microsynth AG, Switzerland) and all final plasmids were verified by analytical restriction digests and partial sequencing.

### Yeast strain construction

All strains are derived from BY4741 and BY4742 (Euroscarf, Germany) [68]. All strains used in this study are summarized in **Supplementary Table 3**. Transformations were performed with the standard lithium acetate method [69] and selection was performed on appropriate selection plates. For genomic integration, plasmids were digested with NotI (NEB) before transformation. Diploid strains were generated by mating and selection by growth on SD plates lacking both L-Lysine and L-Methionine.

### Media and growth conditions

All experiments were performed in synthetic medium (SD; LOFLO yeast nitrogen (ForMedium), 5 g/L ammonium sulfate, 2% glucose, pH was adjusted to 6.0). All experiments were performed in 25 ml glass centrifuge tubes (Schott 2160114, Duran) stirred with 3 × 8 mm magnetic stir bars (Huberlab) using a setup comprised of a water bath (ED (v.2) THERM60, Julabo) set to 30 °C, a multi position magnetic stirrer (Telesystem 15, Thermo Scientific) set to 900 rpm, a 3D printed, custom-made 15-tube holder, and custom-made LED pads (460 nm peak wavelength) located underneath the culture tubes. A white diffusion filter (LEE Filters) was placed between the LED and the culture tube to allow for even illumination. LED intensity was measured at 4 cm distance from the light source using a NOVA power meter and a PD300 photodiode sensor (Ophir Optronics).

### Flow cytometry

All flow cytometry experiments were performed as follows. Cultures were grown overnight starting from single yeast colonies or glycerol stocks, subcultured in fresh medium and grown for at least 16 h in the dark while maintaining an optical density at 600 nm (OD600) lower than 0.4. At the start of the experiment, cells were diluted to an OD600 of 0.01 in 4 ml of medium. Before measurement, cell samples were incubated in SD with 0.1 mg/ml cycloheximide for 3 h at 30 °C to ensure full fluorescent protein maturation. Samples were analyzed using a Cytoflex S flow cytometer (Beckman Coulter). To measure RFP fluorescence, a 488 nm excitation laser and a 625/40 nm emission filter, for YFP, a 488 nm excitation laser and a 525/40 nm emission filter, and for CFP, a 405 nm excitation laser and a 450/45 nm emission filter were used. Data was compensated for fluorescence spill-over and analyzed using R (3.3.2) with the flowCore package [70]. Cells were gated based on forward and side scatter to remove debris and cell aggregates [71]. The data represents the mean fluorescence of analyzed single cells. Strong outliers were removed from the data as follows: First, the fluorescence values were log-transformed. Outliers were defined as data-points with an absolute deviation from the fluorescence distribution median of greater than 4-fold of the median absolute deviation. For the analysis of channel capacity (below), fluorescent levels were normalized by forward-scatter area which functions as a proxy for cell size. At least 5000 cells and typically 20,000 cells were analyzed per experimental condition.

### Information theoretic analysis of flow cytometry data

To quantify the fidelity of information transmission in cellular signaling and gene expression, it is helpful to view the cell as a system in which an input X is mapped to an output Y. Given cellular variability, e.g. due to noise in gene expression, the response of single cells to a particular input signal x_i_ is probabilistic and we can assume the population response to follow a conditional probability distribution P(Y|X=x_i_). The degree to which single cells can distinguish distinct input signals depends on overlap of the respective conditional output distributions. The response fidelity can be assessed using the mutual information (MI) which quantifies the amount of information (in bits) that is gained about one random variable from observing a second random variable.

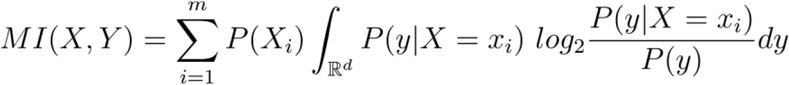

The input distribution P(X), can be seen as the frequency at which inputs (x_1_, x_2_,…, x_m_) occur in the cellular environment. Importantly, the mutual information depends on P(X). To circumvent selection of an arbitrary distribution, the input distribution that maximizes the mutual information can be found. This maximal mutual information is known as the channel capacity (C) of a communication channel.

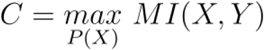

The value 2^C^ can be interpreted as the maximal number of distinct input signals that can possibly be distinguished by the system, in our case the gene expression demultiplexer in single cells, without error. To estimate this channel capacity we make use of a previously described algorithm called SLEMI for “statistical learning based estimation of mutual information” which is implemented as an R-package (see [52] for details on the algorithm). For each input type, we estimate the channel capacity using 10-11 distinct input signals and a two-dimensional output, namely RFP and YFP fluorescence at steady state (mean dose-response data for the analyzed experiments is shown in Fig. 5b-d). We further perform the analysis in the case of using all three input types, meaning 29 input signals after removing redundant signals (no or maximal light exposure).

### Modeling and parameter fitting

The ODE model for the EL222-based expression system was introduced in [25]. The model consists of the following three ordinary differential equations describing Msn2AD-EL222 activation (1), Msn2AD-EL222 dependent mRNA expression (2), and protein expression (3):

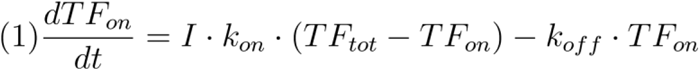

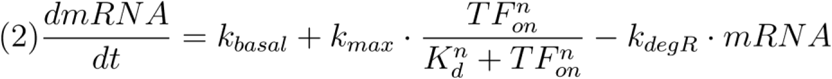

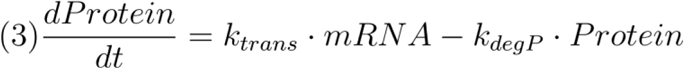

To describe gene expression repression by Mig1RD-EL222, we use a hill-type inhibition in the transcription rate (4). Mig1RD-EL222 activation and downstream protein expression are described by (1) and (2), respectively.

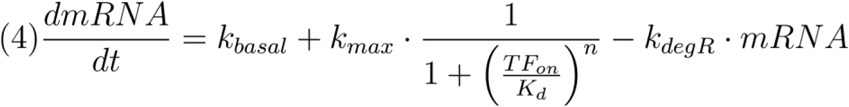

To model transcriptional regulation of a target gene by both Msn2AD-EL222 (denoted as Act) and Mig1RD-EL222 (denoted as Rep) in the diamond-IFFL, we use a multiplication of hill-type activation and inhibition (5).

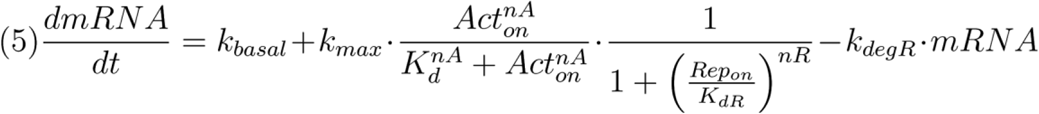

Simulations and model fitting were performed using Matlab (R2014a, Mathworks). The simple expression model (Eq. (1) - (3)) contains 10 free parameters, whereby two of them (kon and koff) are specific to different EL222 mutants while the others are shared. Based on our results from a comparable EL222-based expression system using the same fluorescence reporter gene [25], we fixed parameters values of wt-EL222 koff, TFtot, kdegR, and kdegP (**Supplementary Table 4**). For simplicity, we assume a direct relationship between mean cellular fluorescence measured by flow cytometry and protein expression in the model for parameter fitting. Parameter fitting was performed using a simplex-based search (Nelder-Mead algorithm, “fminsearch” function in Matlab) to minimize the sum of squared residuals (SSR) between the model and the data. We first fitted all model parameters to the data from the wt-EL222 characterization experiments (**Fig. 2c,d**) using different initial parameter values. We then used parameter values with an SSR under a defined threshold and performed a second fit of the EL222-variant dependent parameters kon and koff for the A79Q and AQTrip variants (**Fig. 2c,d**). Parameter values that resulted in the minimal total SSR for all characterization experiments / EL222-variants were used in this study and are summarized in **Supplementary Table 4**. Parameter values for Mig1RD-EL222-mediated repression were obtained by fitting a model based on equations (1),(3), and (4) to the characterization experiment shown in **Fig. 2f** as described above. Parameters describing Mig1RD-EL222 activation, mRNA degradation, and protein expression were fixed to the values obtained from the Msn2AD-EL222 characterization experiments.

### Microscopy Setup

All images were taken with a Nikon Ti-Eclipse inverted microscope (Nikon Instruments), equipped with a 40x, oil-immersion objective (MRH01401, Nikon AG, Egg, Switzerland), Spectra X Light Engine fluorescence excitation light source (Lumencor, USA), pE-100 brightfield light source (CoolLED Ltd., UK), and CMOS camera ORCA-Flash4.0 (Hamamatsu Photonic, Switzerland). The microscope is surrounded by an environmental box (Life Imaging Services, Switzerland) heated to 30 °C. The microscope was operated using NIS-Elements software. A green interference filter was placed into the bright-field light path to avoid activation of optogenetic tools by white light. Z-stacks consisting of 7 images with a step size of 0.4 µm were taken for brightfield and RFP (Excitation line from light source: 550/15 nm, Additional excitation filter: 561/4 nm, beam splitter: HC-BS573, Emission filter: 605/40nm).

### Microscopy sample preparation

Microscopy experiments were performed in 96-square-well glass-bottom plates (MatTek, USA) coated with Concanavalin A using a 2 mg/ml solution (Sigma-Aldrich, USA). Overnight yeast cultures were diluted to an OD600 of 0.05 and grown at 30 °C for 3 h. 200 µl of liquid culture were transferred into coated well plates preheated to 30 °C and cells were allowed to settle for 15 min. The media was removed and wells were washed one time with preheated media and 200 µl of fresh media were added to the well. Experiments were started between 15 and 30 min after washing.

### Microscopy image analysis

The image analysis procedure was performed using custom Matlab scripts and consists of three steps: segmenting individual nuclei (based on nuclear fluorescence), locating fluorescent spots in the nuclear regions, and quantifying the intensity of these spots.

Maximal intensity z-projections were used for RFP images. Background was subtracted from RFP images and images were flatfield corrected based on RFP images taken from wells that only contained media. Nuclei were first enhanced by using the difference of Gaussians algorithm. Nuclear regions were then segmented by manually optimized thresholding. Detected regions that were too big or small to represent nuclei were removed. For each nuclear region, a Difference of Gaussian algorithm was used to enhance spots in the RFP images and spots were identified using thresholding. In order to quantify the intensity of the nuclear spots, the sum of a two-dimensional Gaussian function and a 2D-plane was fitted in a square area around the identified spot with an edge length of 19 pixels [30]. Spot intensity was then defined as the integral of the Gaussian function. For each nucleus / cell, the spot with the highest intensity was defined as the transcription site and spot fluorescence was set to 0 if no spots were detected.

### Carotenoid pathway experiments

For experiments involving carotenoid production, yeast overnight cultures were first diluted to an OD of 0.1 and grown for 6 h. At the start of the experiment, yeast cells were diluted in 4 ml of fresh culture media to an OD600 of 0.2. For PWM experiments cells were first grown in darkness for 2 h and then illuminated for 12 h. For pulse trains experiments illumination was started immediately after inoculation and was performed for 16 h. At the end of the experiment, 800 µl of culture media was sampled and cells were pelleted by centrifugation at 12,000 g and cells were resuspended in 40 µl phosphate buffered saline. To evaluate yeast cell color, 10 µl of cell suspension were spotted on a Immobilon-FL PVDF transfer membrane (Sigma-Aldrich, USA) and dried at 60 °C. Images were taken using a Canon PowerShot SX540HS digital camera using a custom built stand under identical lighting conditions. Measurements of cellular fluorescence were performed using flow cytometry as described in the methods above (“Flow cytometry”).

### Data Availability

Data, plasmids, strains are available from the corresponding author upon request.

### Code Availability

Custom computer code is available from the corresponding author upon request.

## Notes

### Competing Interest Statement

The authors have declared no competing interest.

