## Supplementary Material for "Synthetic gene networks recapitulate dynamic signal decoding and differential gene expression"

### Supplementary Information

Contains:
Supplementary Figure 1
Supplementary Tables 1-4
Supplementary References

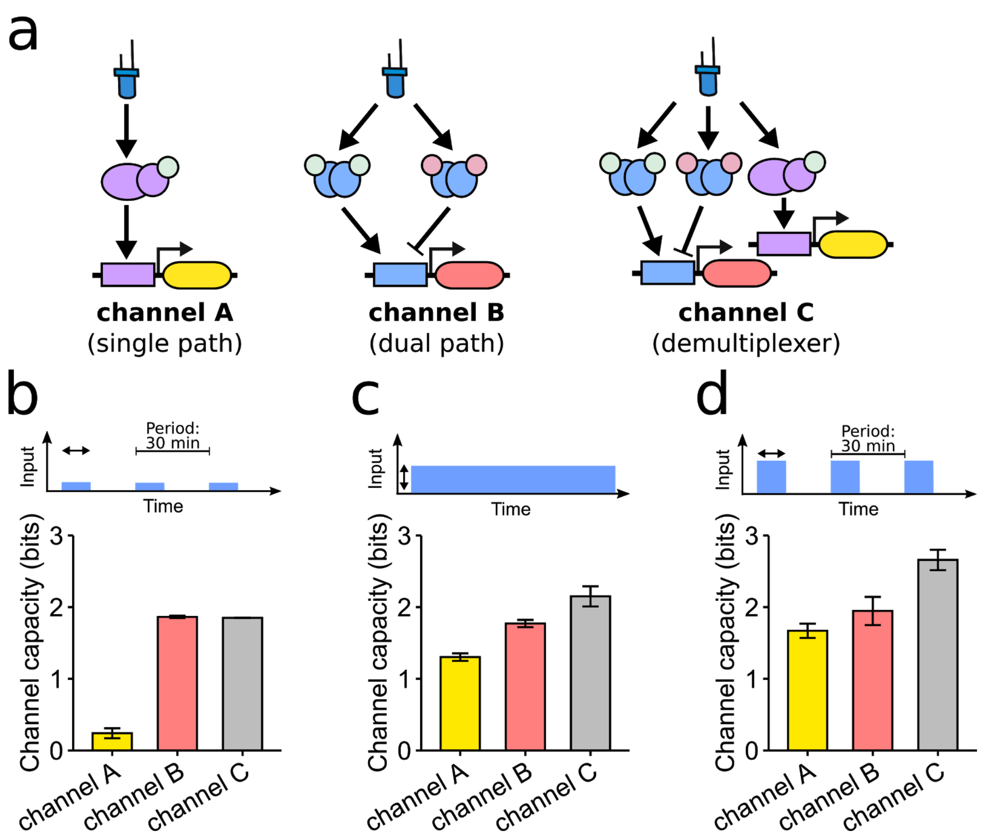


**Supplementary Figure 1. Additional information theoretic analysis of the gene expression demultiplexer. (a)** Schematic of different channels for which the channel capacity was estimated. **(b-d)** Channel capacity was estimated for the three channels shown in (a) using either low intensity PWM (b), AM (c), or high intensity PWM (d) signals as inputs (see Methods for details on analysis). The demultiplexer (channel C) shows an increased channel capacity for AM and high intensity PWM compared to channel A and B. This is not the case for low intensity PWM because there is only a minor response of the branch corresponding to channel A under these conditions (Fig. 5c). Data is taken from experiments shown in Fig 5. Results represent the mean and s.d. calculated based on two independent experiments.

**Supplementary Table 1.** YTK MoClo part sequences used in this study. Coding sequences are capitalized.

| Insert Name | Part Type | Sequence | Source |
|---|---|---|---|
| **Msn2AD** | 3a | gcatcgtctcatcggtctcatatgggccctaaaaagaagcgtaaagtcACGGTCGACCATGATTTCAATAGCGAAGATATTTTATTCCCCATAGA<br>AAGCATGAGTAGTATACAATACGTGGAGAATAATAACCCAAATAATATTAACAACGATGTTATCCCGTATTCTCTAGATATCAA<br>AAACACTGTCTTAGATAGTGCGGATCTCAATGACATTCAAAATCAAGAAACTTCACTGAATTTGGGGCTTCCTCCACTATCTTTC<br>GACTCTCCACTGCCCGTAACGGAAACGATACCATCCACTACCGATAACAGCTTGCATTTGAAAGCTGATAGCAACAAAAATCGC<br>GATGCAAGAACTATTGAAAATGATAGTGAAATTAAGAGTACTAATAATGCTAGTGGCTCTGGGGCAAATCAATACACAACTCT<br>TACTTCACCTTATCCTATGAACGACATTTTGTACAACATGAACAATCCGTTACAATCACCGTCACCTTCATCGGTACCTCAAAATC<br>CGACTATAAATCCTCCCATAAATACAGCAAGTAACGAAACTAATTTATCGCCTCAAACTTCAAATGGTAATGAAACTCTTATATC<br>TCCTCGAGCCCAACAACATACGTCCATTAAAGATAATCGTCTGTCCTTACCTAATGGTGCTAATTCGAATCTTTTCATTGACACT<br>AACCCAAACAATTTGAACGAAAAACTAAGAAATCAATTGAACTCAGATACAAATTCATATTCTAACTCCATTTCTAATTCAAACT<br>CCAATTCTACGGGTAATTTAAATTCCAGTTATTTTAATTCACTGAACATAGACTCCATGCTAGATGATTACGTTTCTAGTGATCTC<br>TTATTGAATGATGATGATGATGACACTAATTTATCACGCCGAAGATTTAGCGACGTTATAACAAACagtgctggtagtgctggttcttga<br>gacctgagacggcat | BY4741 genome |
| **EL222(WT)** | 3b | GAATTCGGGGCAGACGACACACGCGTTGAGGTGCAACCGCCGGCGCAGTGGGTCCTCGACCTGATCGAGGCCAGCCCGATC<br>GCATCGGTCGTGTCCGATCCGCGACTCGCCGACAATCCGCTGATCGCCATCAACCAGGCCTTCACCGACCTGACCGGCTATTCC<br>GAAGAAGAATGCGTCGGCCGCAATTGCCGATTCCTGGCAGGTTCCGGCACCGAGCCGTGGCTGACCGACAAGATCCGCCAAG<br>GCGTGCGCGAGCACAAGCCGGTGCTGGTCGAGATCCTGAACTACAAGAAGGACGGCACGCCGTTCCGCAATGCCGTGCTCGT<br>TGCACCGATCTACGATGACGACGACGAGCTTCTCTATTTCCTCGGCAGCCAGGTCGAAGTCGACGACGACCAGCCCAACATGG<br>GCATGGCGCGCCGCGAACGCGCCGCGGAAATGCTCAAGACGCTGTCGCCGCGCCAGCTCGAGGTTACGACGCTGGTGGCATC<br>GGGCTTGCGCAACAAGGAAGTGGCGGCCCGGCTCGGCCTGTCGGAGAAAACCGTCAAGATGCACCGCGGGCTGGTGATGGA<br>AAAGCTCAACCTGAAGACCAGTGCCGATCTGGTGCGCATTGCCGTCGAAGCCGGAATCTAA | [1] |
| **EL222(A79Q)** | 3b | gctggtGAATTCGGGGCAGACGACACACGCGTTGAGGTGCAACCGCCGGCGCAGTGGGTCCTCGACCTGATCGAGGCCAGCCC<br>GATCGCATCGGTCGTGTCCGATCCGCGACTCGCCGACAATCCGCTGATCGCCATCAACCAGGCCTTCACCGACCTGACCGGCT<br>ATTCCGAAGAAGAATGCGTCGGCCGCAATTGCCGATTCCTGCAAGGTTCCGGCACCGAGCCGTGGCTGACCGACAAGATCCG<br>CCAAGGCGTGCGCGAGCACAAGCCGGTGCTGGTCGAGATCCTGAACTACAAGAAGGACGGCACGCCGTTCCGCAATGCCGTG<br>CTCGTTGCACCGATCTACGATGACGACGACGAGCTTCTCTATTTCCTCGGCAGCCAGGTCGAAGTCGACGACGACCAGCCCAA<br>CATGGGCATGGCGCGCCGCGAACGCGCCGCGGAAATGCTCAAGACGCTGTCGCCGCGCCAGCTCGAGGTTACGACGCTGGT<br>GGCATCGGGCTTGCGCAACAAGGAAGTGGCGGCCCGGCTCGGCCTGTCGGAGAAAACCGTCAAGATGCACCGCGGGCTGGT<br>GATGGAAAAGCTCAACCTGAAGACCAGTGCCGATCTGGTGCGCATTGCCGTCGAAGCCGGAATCTAA | gBlock(IDT) |
| **EL222(AQTrip)** | 3b | gctggtGAATTCGGGGCAGACGACACACGCGTTGAGGTGCAACCGCCGGCGCAGTGGGTCCTCGACCTGATCGAGGCCAGCCC<br>GATCGCATCGATTGTGTCCGATCCGCGACTCGCCGACAATCCGATTATCGCCATCAACCAGGCCTTCACCGACCTGACCGGCTA<br>TTCCGAAGAAGAATGCGTCGGCCGCAATTGCCGATTCCTGCAAGGTTCCGGCACCGAGCCGTGGCTGACCGACAAGATCCGC<br>CAAGGCGTGCGCGAGCACAAGCCGGTGCTGGTCGAGATCCTGAACTACAAGAAGGACGGCACGCCGTTCCGCAATGCCGTGC<br>TCATTGCACCGATCTACGATGACGACGACGAGCTTCTCTATTTCCTCGGCAGCCAGGTCGAAGTCGACGACGACCAGCCCAAC<br>ATGGGCATGGCGCGCCGCGAACGCGCCGCGGAAATGCTCAAGACGCTGTCGCCGCGCCAGCTCGAGGTTACGACGCTGGTG<br>GCATCGGGCTTGCGCAACAAGGAAGTGGCGGCCCGGCTCGGCCTGTCGGAGAAAACCGTCAAGATGCACCGCGGGCTGGTG<br>ATGGAAAAGCTCAACCTGAAGACCAGTGCCGATCTGGTGCGCATTGCCGTCGAAGCCGGAATCTAA | gBlock(IDT) |
| **Mig1RD** | 3a | ATGGGCCCTAAAAAGAAGCGTAAAGTCGATTCACAAGTTCAAGAACTGGAAACATTACCACCCATAAGAAGTTTACCGTTGCC<br>CTTCCCACACATGGACagtgctggtagtgctgg | BY4741 genome |
| **VP16AD-CIB** | 3 | ATGGGCCCTAAAAAGAAGCGTAAAGTCGCCCCCCCGACCGATGTCAGCCTGGGGGACGAGCTCCACTTAGACGGCGAGGAC<br>GTGGCGATGGCGCATGCCGACGCGCTAGACGATTTCGATCTGGACATGTTGGGGGACGGGGATTCCCCGGGTCCGGGATTTA<br>CCCCCCACGACTCCGCCCCCTACGGCGCTCTGGATATGGCCGACTTCGAGTTTGAGCAGATGTTTACCGATGCCCTTGGAATTG<br>ACGAGTACGGTGGGGAATTCCCGGGGATCCGTCGACGCGTCTGCAGAAGCTTCGGCGGCAGCATGAATGGAGCTATAGGAG<br>GTGACCTTTTGCTCAATTTTCCTGACATGTCGGTCCTAGAGCGCCAAAGGGCTCACCTCAAGTACCTCAATCCCACCTTTGATTC<br>TCCTCTCGCCGGCTTCTTTGCCGATTCTTCAATGATTACCGGCGGCGAGATGGACAGCTATCTTTCGACTGCCGGTTTGAATCTT<br>CCGATGATGTACGGTGAAACGACGGTGGAAGGTGATTCAAGACTCTCAATTTCGCCGGAAACGACGCTTGGGACTGGAAATT<br>TCAAGAAACGGAAGTTTGATACAGAGACTAAGGATTGTAATGAGAAGAAGAAGAAGATGACGATGAACAGAGATGACCTAG<br>TAGAAGAAGGAGAAGAAGAGAAGTCGAAAATAACAGAGCAAAACAATGGGAGCACAAAAAGCATCAAGAAGATGAAACAC<br>AAAGCCAAGAAAGAAGAGAACAATTTCTCTAATGATTCATCTAAAGTGACGAAGGAATTGGAGAAAACGGATTATATTCATGT<br>TCGTGCACGACGAGGCCAAGCCACTGATAGTCACAGCATAGCAGAACGAGTTAGAAGAGAAAAGATCAGTGAGAGAATGAA<br>GTTTCTACAAGATTTGGTTCCTGGATGCGACAAGATCACAGGCAAAGCAGGGATGCTTGATGAAATCATTAACTATGTTCAGT<br>CTCTTCAGAGACAAATCGAGTTCTTATCGATGAAACTAGCAATTGTGAATCCAAGGCCGGATTTTGATATGGATGACATTTTTG<br>CCAAAGAGGTTGCCTCAACTCCAATGACTGTGGTGCCATCTCCTGAAATGGTTCTTTCCGGTTATTCTCATGAGATGGTTCACTC<br>TGGTTATTCTAGTGAGATGGTTAACTCCGGTTACCTTCATGTCAATCCAATGCAGCAAGTGAATACCAGTTCTGATCCATTGTCA | [2] |

| | | | |
|---|---|---|---|
| | | TGCTTCAACAATGGCGAAGCTCCTTCGATGTGGGACTCTCATGTGCAGAATCTCTATGGCAATTTAGGAGTACCGGTCATCGA<br>GCTCGAGCTGCAGATGAATCGTAGATACTGA | |
| **LexA-CRY2PHR(W349R)** | 3 | ATGAAAGCGTTAACGGCCAGGCAACAAGAGGTGTTTGATCTCATCCGTGATCACATCAGCCAGACAGGTATGCCGCCGACGC<br>GTGCGGAAATCGCGCAGCGTTTGGGGTTCCGTTCCCCAAACGCGGCTGAAGAACATCTGAAGGCGCTGGCACGCAAAGGCGT<br>TATTGAAATTGTTTCCGGCGCATCACGCGGGATTCGTCTGTTGCAGGAAGAGGAAGAAGGGTTGCCGCTGGTAGGTCGTGTG<br>GCTGCCGGTGAACCACTTCTGGCGCAACAGCATATTGAAGGTCATTATCAGGTCGATCCTTCCTTATTCAAGCCGAATGCTGAT<br>TTCCTGCTGCGCGTCAGCGGGATGTCGATGAAAGATATCGGCATTATGGATGGTGACTTGCTGGCAGTGCATAAAACTCAGGA<br>TGTACGTAACGGTCAGGTCGTTGTCGCACGTATTGATGACGAAGTTACCGTTAAGCGCCTGAAAAAACAGGGCAATAAAGTC<br>GAACTGTTGCCAGAAAATAGCGAGTTTAAACCAATTGTCGTAGATCTTCGTCAGCAGAGCTTCACCATTGAAGGGCTGGCGGT<br>TGGGGTTATTCGCAACGGCGACTGGCTGGAATTCCCGGGGATCCGTCGACCATGGCGACCGCTCGAGTCGACCTGCAGCCAA<br>GCTAATTCCGGGCGAATTTCTTATGATTTAGGCGGCAGCATGAAGATGGACAAAAAGACTATAGTTTGGTTTAGAAGGGACCT<br>AAGGATTGAGGATAATCCTGCATTAGCAGCAGCTGCTCACGAAGGATCTGTTTTTCCTGTCTTCATTTGGTGTCCTGAAGAAGA<br>AGGACAGTTTTATCCTGGAAGAGCTTCAAGATGGTGGATGAAACAATCACTTGCTCACTTATCTCAATCCTTGAAGGCTCTTGG<br>ATCTGACCTCACTTTAATCAAAACCCACAACACGATTTCAGCGATCTTGGATTGTATCCGCGTTACCGGTGCTACAAAAGTCGTC<br>TTTAACCACCTCTATGATCCTGTTTCGTTAGTTCGGGACCATACCGTAAAGGAGAAGCTGGTGGAACGTGGGATCTCTGTGCA<br>AAGCTACAATGGAGATCTATTGTATGAACCGTGGGAGATATACTGCGAAAAGGGCAAACCTTTTACGAGTTTCAATTCTTACT<br>GGAAGAAATGCTTAGATATGTCGATTGAATCCGTTATGCTTCCTCCTCCTTGGCGGTTGATGCCAATAACTGCAGCGGCTGAA<br>GCGATTTGGGCGTGTTCGATTGAAGAACTAGGGCTGGAGAATGAGGCCGAGAAACCGAGCAATGCGTTGTTAACTAGAGCTT<br>GGTCCCCAGGATGGAGCAATGCTGATAAGTTACTAAATGAGTTCATCGAGAAGCAGTTGATAGATTATGCAAAGAACAGCAA<br>GAAAGTTGTTGGGAATTCTACTTCACTACTTTCTCCGTATCTCCATTTCGGGGAAATAAGCGTCAGACACGTTTTCCAGTGTGCC<br>CGGATGAAACAAATTATATGGGCAAGAGATAAGAACAGTGAAGGAGAAGAAAGTGCAGATCTTTTTCTTAGGGGAATCGGTT<br>TAAGAGAGTATTCTCGGTATATATGTTTCAACTTCCCGTTTACTCACGAGCAATCGTTGTTGAGTCATCTTCGGTTTTTCCCTTGG<br>GATGCTGATGTTGATAAGTTCAAGGCCTGGAGACAAGGCAGGACCGGTTATCCGTTGGTGGATGCCGGAATGAGAGAGCTTA<br>GGGCTACCGGATGGATGCATAACAGAATAAGAGTGATTGTTTCAAGCTTTGCTGTGAAGTTTCTTCTCCTTCCATGGAAATGG<br>GGAATGAAGTATTTCTGGGATACACTTTTGGATGCTGATTTGGAATGTGACATCCTTGGCTGGCAGTATATCTCTGGGAGTATC<br>CCCGATGGCCACGAGCTTGATCGCTTGGACAATCCCGCGTTACAAGGCGCCAAATATGACCCAGAAGGTGAGTACATAAGGC<br>AATGGCTTCCCGAGCTTGCGAGATTGCCAACTGAATGGATCCATCATCCATGGGACGCTCCTTTAACCGTACTCAAAGCTTCTG<br>GTGTGGAACTCGGAACAAACTATGCGAAACCCATTGTAGACATCGACACAGCTCGTGAGCTACTAGCTAAAGCTATTTCAAGA<br>ACCCGTGAAGCACAGATCATGATCGGAGCAGCAGCCCGGGTCGACCTGCAGCCAAGCTAA | [3] |
| **prLexACYCmin** | 2 | AACGggccgcaataatatataaacctgtataatataaccttgaagactatatttcttttcgagctccctaggtgctgtatatactcacagcataactgtatatacaccc<br>agggtctaggtgctgtatatactcacagcataactgtatatacacccagggtctaggtgctgtatatactcacagcataactgtatatacacccagggtctaggtgctgt<br>atatactcacagcataactgtatatacacccagggtctagagcatgtgctctgtatgtatataaaactcttgttttcttcttttctcttctttccttatacattaggacctttg<br>cagcataaattactatacttctatactagtggatcccccgggctgcaggaattcgatatcaagctTATG | [4] |

| | | | |
|---|---|---|---|
| **dCas9** | 3a | tatgAGTGATAAAAAGTATTCTATTGGTTTAGCCATCGGCACTAATTCCGTTGGATGGGCTGTCATAACCGATGAATACAAAGTA<br>CCTTCAAAGAAATTTAAGGTGTTGGGGAACACAGACCGTCATTCGATTAAAAAGAATCTTATCGGTGCCCTCCTATTCGATAGT<br>GGCGAAACGGCAGAGGCGACTCGCCTGAAACGAACCGCTCGGAGAAGGTATACACGTCGCAAGAACCGAATATGTTACTTAC<br>AAGAAATTTTTAGCAATGAGATGGCCAAAGTTGACGATTCTTTCTTTCACCGTTTGGAAGAGTCCTTCCTTGTCGAAGAGGACA<br>AGAAACATGAACGGCACCCCATCTTTGGAAACATAGTAGATGAGGTGGCATATCATGAAAAGTACCCAACGATTTATCACCTC<br>AGAAAAAAGCTAGTTGACTCAACTGATAAAGCGGACCTGAGGTTAATCTACTTGGCTCTTGCCCATATGATAAAGTTCCGTGG<br>GCACTTTCTCATTGAGGGTGATCTAAATCCGGACAACTCGGATGTCGACAAACTGTTCATCCAGTTAGTACAAACCTATAATCA<br>GTTGTTTGAAGAGAACCCTATAAATGCAAGTGGCGTGGATGCGAAGGCTATTCTTAGCGCCCGCCTCTCTAAATCCCGACGGC<br>TAGAAAACCTGATCGCACAATTACCCGGAGAGAAGAAAAATGGGTTGTTCGGTAACCTTATAGCGCTCTCACTAGGCCTGACA<br>CCAAATTTTAAGTCGAACTTCGACTTAGCTGAAGATGCCAAATTGCAGCTTAGTAAGGACACGTACGATGACGATCTCGACAA<br>TCTACTGGCACAAATTGGAGATCAGTATGCGGACTTATTTTTGGCTGCCAAAAACCTTAGCGATGCAATCCTCCTATCTGACAT<br>ACTGAGAGTTAATACTGAGATTACCAAGGCGCCGTTATCCGCTTCAATGATCAAAAGGTACGATGAACATCACCAAGACTTGA<br>CACTTCTCAAGGCCCTAGTCCGTCAGCAACTGCCTGAGAAATATAAGGAAATATTCTTTGATCAGTCGAAAAACGGGTACGCA<br>GGTTATATTGACGGCGGAGCGAGTCAAGAGGAATTCTACAAGTTTATCAAACCCATATTAGAGAAGATGGATGGGACGGAAG<br>AGTTGCTTGTAAAACTCAATCGCGAAGATCTACTGCGAAAGCAGCGGACTTTCGACAACGGTAGCATTCCACATCAAATCCACT<br>TAGGCGAATTGCATGCTATACTTAGAAGGCAGGAGGATTTTTATCCGTTCCTCAAAGACAATCGTGAAAAGATTGAGAAAATC<br>CTAACCTTTCGCATACCTTACTATGTGGGACCCCTGGCCCGAGGGAACTCTCGGTTCGCATGGATGACAAGAAAGTCCGAAGA<br>AACGATTACTCCATGGAATTTTGAGGAAGTTGTCGATAAAGGTGCGTCAGCTCAATCGTTCATCGAGAGGATGACCAACTTTG<br>ACAAGAATTTACCGAACGAAAAAGTATTGCCTAAGCACAGTTTACTTTACGAGTATTTCACAGTGTACAATGAACTCACGAAAG<br>TTAAGTATGTCACTGAGGGCATGCGTAAACCCGCCTTTCTAAGCGGAGAACAGAAGAAAGCAATAGTAGATCTGTTATTCAAG<br>ACCAACCGCAAAGTGACAGTTAAGCAATTGAAAGAGGACTACTTTAAGAAAATTGAATGCTTCGATTCTGTCGAGATCTCCGG<br>GGTAGAAGATCGATTTAATGCGTCACTTGGTACGTATCATGACCTCCTAAAGATAATTAAAGATAAGGACTTCCTGGATAACG<br>AAGAGAATGAAGATATCTTAGAAGATATAGTGTTGACTCTTACCCTCTTTGAAGATCGGGAAATGATTGAGGAAAGACTAAAA<br>ACATACGCTCACCTGTTCGACGATAAGGTTATGAAACAGTTAAAGAGGCGTCGCTATACGGGCTGGGGACGATTGTCGCGGA<br>AACTTATCAACGGGATAAGAGACAAGCAAAGTGGTAAAACTATTCTCGATTTTCTAAAGAGCGACGGCTTCGCCAATAGGAAC<br>TTTATGCAGCTGATCCATGATGACTCTTTAACCTTCAAAGAGGATATACAAAAGGCACAGGTTTCCGGACAAGGGGACTCATT<br>GCACGAACATATTGCGAATCTTGCTGGTTCGCCAGCCATCAAAAAGGGCATACTCCAGACAGTCAAAGTAGTGGATGAGCTAG<br>TTAAGGTCATGGGACGTCACAAACCGGAAAACATTGTAATCGAGATGGCACGCGAAAATCAAACGACTCAGAAGGGGCAAAA<br>AAACAGTCGAGAGCGGATGAAGAGAATAGAAGAGGGTATTAAAGAACTGGGCAGCCAGATCTTAAAGGAGCATCCTGTGGA<br>AAATACCCAATTGCAGAACGAGAAACTTTACCTCTATTACCTACAAAATGGAAGGGACATGTATGTTGATCAGGAACTGGACA<br>TAAACCGTTTATCTGATTACGACGTCGATGCCATTGTACCCCAATCCTTTTTGAAGGACGATTCAATCGACAATAAAGTGCTTAC<br>ACGCTCGGATAAGAACCGAGGGAAAAGTGACAATGTTCCAAGCGAGGAAGTCGTAAAGAAAATGAAGAACTATTGGCGGCA<br>GCTCCTAAATGCGAAACTGATAACGCAAAGAAAGTTCGATAACTTAACTAAAGCTGAGAGGGGTGGCTTGTCTGAACTTGACA<br>AGGCCGGATTTATTAAACGTCAGCTCGTGGAAACCCGCCAAATCACAAAGCATGTTGCACAGATACTAGATTCCCGAATGAAT<br>ACGAAATACGACGAGAACGATAAGCTGATTCGGGAAGTCAAAGTAATCACTTTAAAGTCAAAATTGGTGTCGGACTTCAGAA<br>AGGATTTTCAATTCTATAAAGTTAGGGAGATAAATAACTACCACCATGCGCACGACGCTTATCTTAATGCCGTCGTAGGGACCG<br>CACTCATTAAGAAATACCCGAAGCTAGAAAGTGAGTTTGTGTATGGTGATTACAAAGTTTATGACGTCCGTAAGATGATCGCG<br>AAAAGCGAACAGGAGATAGGCAAGGCTACAGCCAAATACTTCTTTTATTCTAACATTATGAATTTCTTTAAGACGGAAATCACT<br>CTGGCAAACGGAGAGATACGCAAACGACCTTTAATTGAAACCAATGGGGAGACAGGTGAAATCGTATGGGATAAGGGCCGG<br>GACTTCGCGACGGTGAGAAAAGTTTTGTCCATGCCCCAAGTCAACATAGTAAAGAAAACTGAGGTGCAGACCGGAGGGTTTT<br>CAAAGGAATCGATTCTTCCAAAAAGGAATAGTGATAAGCTCATCGCTCGTAAAAAGGACTGGGACCCGAAAAAGTACGGTGG<br>CTTCGATAGCCCTACAGTTGCCTATTCTGTCCTAGTAGTGGCAAAAGTTGAGAAGGGAAAATCCAAGAAACTGAAGTCAGTCA<br>AAGAATTATTGGGGATAACGATTATGGAGCGCTCGTCTTTTGAAAAGAACCCCATCGACTTCCTTGAGGCGAAAGGTTACAAG<br>GAAGTAAAAAAGGATCTCATAATTAAACTACCAAAGTATAGTCTGTTTGAGTTAGAAAATGGCCGAAAACGGATGTTGGCTAG<br>CGCCGGAGAGCTTCAAAAGGGGAACGAACTCGCACTACCGTCTAAATACGTGAATTTCCTGTATTTAGCGTCCCATTACGAGA<br>AGTTGAAAGGTTCACCTGAAGATAACGAACAGAAGCAACTTTTTGTTGAGCAGCACAAACATTATCTCGACGAAATCATAGAG<br>CAAATTTCGGAATTCAGTAAGAGAGTCATCCTAGCTGATGCCAATCTGGACAAAGTATTAAGCGCATACAACAAGCACAGGGA<br>TAAACCCATACGTGAGCAGGCGGAAAATATTATCCATTTGTTTACTCTTACCAACCTCGGCGCTCCAGCCGCATTCAAGTATTTT<br>GACACAACGATAGATCGCAAACGATACACTTCTACCAAGGAGGTGCTAGACGCGACACTGATTCACCAATCCATCACGGGATT<br>ATATGAAACTCGGATAGATTTGTCACAGCTTGGGGGTGACgg | [5] |
| **Mxi1** | 3b | TTCTGAGGGAGCTCCCAAGAAAAAGCGCAAGGTAGGTAGTTCCAAGCTTGGCGGCAGCGGCGGCAGCATGGAACGTGTGAG<br>AATGATTAATGTGCAAAGGCTGTTAGAAGCCGCAGAGTTTTTAGAAAGAAGAGAAAGAGAATGCGAACACGGGTATGCCAGT<br>TCTTTCCCTAGCATGCCCTCTCCCAGAGGCgg | [5] |
| **VP64** | 3b | TTCTAGCAGGGCTGACCCCAAGAAGAAGAGGAAGGTGGAGGCCAGCGGTTCCGGACGGGCTGACGCATTGGACGATTTTGA<br>TCTGGATATGCTGGGAAGTGACGCCCTCGATGATTTTGACCTTGACATGCTTGGTTCGGATGCCCTTGATGACTTTGACCTCGA<br>CATGCTCGGCAGTGACGCCCTTGATGATTTCGACCTGGACATGCTGATTAACTCTAGAAGTTCCGGATCTCCGAAAAAGAAAC<br>GCAAAGTTgg | [5] |
| **HH-LexABSsgRNA-HDV** | 3 | tatgatacagctgatgagtccgtgaggacgaaacgagtaagctcgtcctgtatatacacccagggtctgttttagagctagaaatagcaagttaaaataaggctagtc<br>cgttatcaacttgaaaaagtggcaccgagtcggtgcttttggccggcatggtcccagcctcctcgctggcgccggctgggcaacatgcttcggcatggcgaatgggac | gBlock(IDT) |
| **HH-GAL7sgRNA-HDV** | 3 | tatgacagttctgatgagtccgtgaggacgaaacgagtaagctcgtcaactgttgaccgtgatccgagttttagagctagaaatagcaagttaaaataaggctagtcc<br>gttatcaacttgaaaaagtggcaccgagtcggtgcttttggccggcatggtcccagcctcctcgctggcgccggctgggcaacatgcttcggcatggcgaatgggac | gBlock(IDT) |
| **pr5xBS-CYC180** | 2 | aacggtcacagcttgtctgtaagcggatgccgggagcagacaagcccgtcagggcgcgtcagcgggtgttggcgggtgtcggggctggcttaactatgcggcatcag<br>agcagattgtactgagagtgcaccatatggacatattgtcgttagaacgcggctacaattaatacataaccttatgtatcatacacatacgatttaggtgacactatag<br>aacggccgccagctgaagcttgcgttcgctactagtanctagnctttagtccatgtctagtanctagcctttagtccatgtctagtanctagcctttagtccatgtctagt<br>agctagcctttagtccatgtctagtagctagcctttagtccatgtctagagtgctgacactacaggcatatatatatgtgtgcgacgacacatgatcatatggcatgcat | [6] |

| | | | |
|---|---|---|---|
| | | gtgctctgtatgtatataaaactcttgttttcttcttttctctaaatattctttccttatacattaggacctttgcagcataaattactatacttctatagacacacaaacaca<br>aatacattattaaaacaa | |
| **prGal7** | 2 | aacggacggtagcaacaagaatatagcacgagccgcggagttcatttcgttacttttgatatcactcacaactattgcgaagcgcttcagtgaaaaaatcataaggaa<br>aagttgtaaatattattggtagtattcgtttggtaaagtagaggggggtaatttttcccctttattttgttcatacattcttaaattgctttgcctctccttttggaaagctata<br>cttcggagcactgttgagcgaaggctcattagatatattttctgtcattttccttaacccaaaaataagggaaagggtccaaaaagcgctcggacaactgttgaccgtg<br>atccgaaggactggctatacagtgttcacaaaatagccaagctgaaaataatgtgtagctatgttcagttagtttggctagcaaagatataaaagcaggtcggaaata<br>tttatgggcattattatgcagagcatcaacatgataaaaaaaaacagttgaatattccctcaaaa | BY4741 genome |
| **ycrtE** | 3 | ATGGACTACGCTAACATCTTGACCGCTATCCCATTGGAGTTCACCCCACAAGACGACATCGTTTTGTTGGAACCATACCACTACT<br>TGGGTAAGAACCCAGGTAAGGAAATCAGATCTCAATTGATCGAGGCTTTCAACTACTGGTTGGACGTTAAGAAGGAAGACTT<br>GGAAGTTATCCAAAACGTTGTTGGTATGTTGCACACCGGTTCTTTGTTGATGGACGACGTTGAAGACTCTTCTGTTTTGAGAAG<br>AGGTTCTCCAGTTGCTCACTTGATCTACGGTATCCCACAAACCATCAACACCGCTAACTACGTTTACTTCTTGGCTTACCAAGAA<br>ATCTTCAAGCTCAGACCAACCCCAATCCCAATGCCAGTTATCCCACCATCTTCTGCTTCTTTGCAATCTTCTGTTTCTTCTGCTTCT<br>TCTTCTTCTTCTGCTTCTTCTGAAAACGGTGGAACCTCTACCCCAAACTCTCAAATCCCATTCTCTAAGGACACCTACTTGGACAA<br>GGTTATCACCGACGAAATCTTGTCTTTGCACAGAGGTCAAGGTTTGGAATTGTTCTGGAGAGACTCTTTGACCTGTCCATCTGA<br>AGAAGAATACGTTAAGATGGTTTTGGGTAAGACCGGTGGTTTGTTCAGAATCGCTGTTAGATTGATGATGGCTAAGTCTGAAT<br>GTGACATCGACTTCGTTCAATTGGTTAACTTGATCTCTATCTACTTCCAAATCCGCGACGACTACATGAACTTGCAATCTTCTGA<br>ATACGCTCACAACAAGAACTTCGCTGAAGACTTGACCGAAGGTAAGTTCTCTTTCCCAACCATCCACTCTATCCACGCTAACCCA<br>TCTAGTAGATTGGTTATCAACACCTTGCAAAAGAAGTCTACCTCTCCAGAAATCTTGCACCACTGTGTTAACTACATGAGAACC<br>GAAACCCACTCTTTCGAATACACCCAAGAAGTTTTGAACACCTTGTCTGGTGCTTTGGAAAGAGAATTGGGTAGATTGCAAGG<br>TGAGTTCGCTGAAGCTAACAGTAGAATGGACTTGGGTGACGTTGACTCTGAAGGTAGAACCGGTAAGAACGTTAAGTTGGAA<br>GCTATCTTGAAGAAGTTGGCTGACATCCCATTGTAG | [7] |
| **ycrtI** | 3 | ATGGGTAAGGAACAAGACCAAGACAAGCCAACCGCTATCATCGTTGGTTGTGGTATCGGTGGTATCGCTACCGCTGCTAGATT<br>GGCTAAGGAAGGTTTCCAAGTTACCGTTTTCGAAAAGAACGACTACTCTGGTGGTAGATGTTCTTTGATCGAACGCGACGGTT<br>ACAGATTCGACCAAGGTCCATCTTTGTTGTTGTTGCCAGACTTGTTCAAGCAAACCTTCGAAGACTTGGGTGAAAAGATGGAA<br>GACTGGGTTGACTTGATCAAGTGTGAACCAAACTACGTTTGTCACTTCCACGACGAAGAAACCTTCACCTTCTCTACCGACATG<br>GCTTTGTTGAAGAGAGAAGTTGAAAGATTCGAAGGTAAGGACGGTTTCGACAGATTCTTGTCTTTCATCCAAGAAGCTCACAG<br>ACACTACGAATTGGCTGTTGTTCACGTTTTGCAAAAGAACTTCCCAGGTTTCGCTGCTTTCTTGAGATTGCAATTCATCGGTCAA<br>ATCTTGGCTTTGCACCCATTCGAATCTATCTGGACCAGAGTTTGTAGATACTTCAAGACCGACAGATTGAGAAGAGTTTTCTCTT<br>TCGCTGTTATGTACATGGGTCAATCTCCATACTCTGCTCCAGGAACCTACTCTTTGTTGCAATACACCGAATTGACCGAAGGTAT<br>CTGGTATCCAAGAGGTGGTTTCTGGCAAGTTCCAAACACCTTGTTGCAAATCGTTAAGAGAAACAACCCATCTGCTAAGTTCAA<br>CTTCAACGCTCCAGTTTCTCAAGTTTTGTTGTCTCCAGCTAAGGACAGAGCTACCGGTGTTAGATTGGAATCTGGTGAAGAACA<br>CCACGCTGACGTTGTTATCGTTAACGCTGACTTGGTTTACGCTTCTGAACACTTGATCCCAGACGACGCTAGAAACAAGATCGG<br>TCAATTGGGTGAAGTTAAGCGATCTTGGTGGGCTGACTTGGTTGGTGGTAAGAAGTTGAAGGGTTCTTGTTCTTCTTTGTCTTT<br>CTACTGGTCTATGGACAGAATCGTTGACGGTTTGGGTGGTCACAACATCTTCTTGGCTGAAGACTTCAAGGGTTCTTTCGACAC<br>CATCTTCGAAGAATTGGGTTTGCCAGCTGACCCATCTTTCTACGTTAACGTTCCAAGTAGAATCGACCCATCTGCTGCTCCAGAA<br>GGTAAGGACGCTATCGTTATCTTGGTTCCATGTGGTCACATCGACGCTTCTAACCCACAAGACTACAACAAGTTGGTTGCTAGA<br>GCTAGAAAGTTCGTTATCCAAACCTTGTCTGCTAAGTTGGGTTTGCCAGACTTCGAAAAGATGATCGTTGCTGAAAAGGTTCAC<br>GACGCTCCATCTTGGGAAAAGGAATTCAACTTGAAGGACGGTTCTATCTTGGGTTTGGCTCACAACTTCATGCAAGTTTTGGGT<br>TTCAGACCATCTACCAGACACCCAAAGTACGACAAGTTGTTCTTCGTTGGTGCTTCTACCCACCCAGGAACCGGTGTTCCAATC<br>GTTTTGGCTGGTGCTAAGTTGACCGCTAACCAAGTTTTGGAATCTTTCGACCGATCTCCAGCTCCAGACCCAAACATGTCTTTGT<br>CTGTTCCATACGGTAAGCCATTGAAGTCTAACGGAACCGGTATCGACTCTCAAGTTCAATTGAAGTTCATGGACTTGGAAAGAT<br>GGGTTTACTTGTTGGTTTTGTTGATCGGTGCTGTTATCGCTCGATCTGTTGGTGTTTTGGCTTTCTAG | [7] |
| **ycrtYB** | 3 | ATGACCGCTTTGGCTTACTACCAAATCCACTTGATCTACACCTTGCCAATCTTGGGTTTGTTGGGTTTGTTGACCTCTCCAATCTT<br>GACCAAGTTCGACATCTACAAAATCTCTATCTTGGTTTTCATCGCTTTCTCTGCTACCACCCCATGGGACTCTTGGATCATCAGA<br>AACGGTGCTTGGACCTACCCATCTGCTGAATCTGGTCAAGGTGTTTTCGGAACCTTCTTGGACGTTCCATACGAAGAATACGCT<br>TTCTTCGTTATCCAAACCGTTATCACAGGTTTGGTTTACGTTTTGGCTACCAGACACTTGTTGCCATCTTTGGCTTTGCCAAAGAC<br>CCGATCTTCTGCTTTGTCTTTGGCTTTGAAGGCTTTGATCCCATTGCCAATCATCTACTTGTTCACCGCTCACCCATCTCCATCTCC<br>AGACCCATTGGTTACCGACCACTACTTCTACATGAGAGCTTTGTCTTTGTTGATCACCCCACCAACCATGTTGTTGGCTGCTTTG<br>TCTGGTGAATACGCTTTCGACTGGAAGTCTGGTAGAGCTAAGTCTACCATCGCTGCTATCATGATCCCAACCGTTTACTTGATCT<br>GGGTTGACTACGTTGCTGTTGGTCAAGACTCTTGGTCTATCAACGACGAAAAGATCGTTGGTTGGAGATTGGGTGGTGTTTTG<br>CCAATCGAAGAAGCTATGTTCTTCTTGTTGACCAACTTGATGATCGTTTTGGGTTTGTCTGCTTGTGACCACACCCAGGCTTTGT<br>ACTTGTTGCACGGTAGAACCATCTACGGTAACAAGAAGATGCCATCTTCTTTCCCATTGATCACCCCACCAGTTTTGTCTTTGTT<br>CTTCTCTAGTAGACCATACTCTTCTCAACCAAAGAGAGACTTGGAATTGGCTGTTAAGTTGTTGGAAGAAAAGTCTCGATCTTT<br>CTTCGTTGCTTCTGCTGGTTTCCCATCTGAAGTTAGAGAAAGATTGGTTGGTTTGTACGCTTTCTGTAGAGTTACCGACGACTTG<br>ATCGACTCTCCAGAAGTTTCTTCTAACCCACACGCTACCATCGACATGGTTTCTGACTTCTTGACCTTGTTGTTCGGTCCACCATT<br>GCACCCATCTCAACCAGACAAAATCTTGTCTTCTCCATTGTTGCCACCATCTCACCCAAGTAGACCAACAGGTATGTACCCATTG<br>CCACCACCACCATCTTTGTCTCCAGCTGAATTGGTTCAATTCTTGACCGAAAGAGTTCCAGTTCAATACCACTTCGCTTTCAGAT<br>TGTTGGCTAAGTTGCAAGGTTTGATCCCAAGATACCCATTGGACGAATTGTTGAGAGGTTACACCACCGACTTGATCTTCCCAT<br>TGTCTACCGAAGCTGTTCAAGCTAGAAAGACCCCAATCGAAACCACCGCTGACTTGTTGGACTACGGTTTGTGTGTTGCTGGTT<br>CTGTTGCTGAATTGTTGGTTTACGTTTCTTGGGCTTCTGCTCCATCTCAAGTTCCAGCTACCATCGAAGAAAGAGAAGCTGTTTT<br>GGTTGCTAGTAGAGAAATGGGAACCGCTTTGCAATTGGTTAACATCGCTAGAGACATCAAGGGTGACGCTACCGAAGGTAGA<br>TTCTACTTGCCATTGTCTTTCTTCGGTTTGCGCGACGAATCTAAGTTGGCTATCCCAACCGACTGGACCGAACCAAGACCACAA<br>GACTTCGACAAGTTGTTGTCTTTGTCTCCATCTTCTACCTTGCCATCTTCTAACGCTTCTGAATCTTTCAGATTCGAATGGAAGAC<br>CTACTCTTTGCCATTGGTTGCTTACGCTGAAGACTTGGCTAAGCACTCTTACAAGGGTATCGACAGATTGCCAACCGAAGTTCA<br>AGCTGGTATGAGAGCTGCTTGTGCTTCTTACTTGTTGATCGGTAGAGAAATCAAGGTTGTTTGGAAGGGTGACGTTGGTGAAA | [7] |

| | | | |
|---|---|---|---|
| | | GAAGAACCGTTGCTGGTTGGAGAAGAGTTAGAAAGGTTTTGTCTGTTGTTATGTCTGGTTGGGAAGGTCAATAG | |
| **tHMG** | 3 | ATGACTGCAGACCAATTGGTGAAGACTGAAGTCACCAAGAAGTCTTTTACTGCTCCTGTACAAAAGGCTTCTACACCAGTTTTA<br>ACCAATAAAACAGTCATTTCTGGATCGAAAGTCAAAAGTTTATCATCTGCGCAATCGAGCTCATCAGGACCTTCATCATCTAGT<br>GAGGAAGATGATTCCCGCGATATTGAAAGCTTGGATAAGAAAATACGTCCTTTAGAAGAATTAGAAGCATTATTAAGTAGTGG<br>AAATACAAAACAATTGAAGAACAAAGAGGTCGCTGCCTTGGTTATTCACGGTAAGTTACCTTTGTACGCTTTGGAGAAAAAAT<br>TAGGTGATACTACGAGAGCGGTTGCGGTACGTAGGAAGGCTCTTTCAATTTTGGCAGAAGCTCCTGTATTAGCATCTGATCGT<br>TTACCATATAAAAATTATGACTACGACCGCGTATTTGGCGCTTGTTGTGAAAATGTTATAGGTTACATGCCTTTGCCCGTTGGT<br>GTTATAGGCCCCTTGGTTATCGATGGTACATCTTATCATATACCAATGGCAACTACAGAGGGTTGTTTGGTAGCTTCTGCCATG<br>CGTGGCTGTAAGGCAATCAATGCTGGCGGTGGTGCAACAACTGTTTTAACTAAGGATGGTATGACAAGAGGCCCAGTAGTCC<br>GTTTCCCAACTTTGAAAAGATCTGGTGCCTGTAAGATATGGTTAGACTCAGAAGAGGGACAAAACGCAATTAAAAAAGCTTTT<br>AACTCTACATCAAGATTTGCACGTCTGCAACATATTCAAACTTGTCTAGCAGGAGATTTACTCTTCATGAGATTTAGAACAACTA<br>CTGGTGACGCAATGGGTATGAATATGATTTCTAAGGGTGTCGAATACTCATTAAAGCAAATGGTAGAAGAGTATGGCTGGGA<br>AGATATGGAGGTTGTCTCCGTTTCTGGTAACTACTGTACCGACAAAAAACCAGCTGCCATCAACTGGATCGAAGGTCGTGGTA<br>AGAGTGTCGTCGCAGAAGCTACTATTCCTGGTGATGTTGTCAGAAAAGTGTTAAAAAGTGATGTTTCCGCATTGGTTGAGTTG<br>AACATTGCTAAGAATTTGGTTGGATCTGCAATGGCTGGGTCTGTTGGTGGATTTAACGCACATGCAGCTAATTTAGTGACAGC<br>TGTTTTCTTGGCATTAGGACAAGATCCTGCACAAAATGTCGAAAGTTCCAACTGTATAACATTGATGAAAGAAGTGGACGGTG<br>ATTTGAGAATTTCCGTATCCATGCCATCCATCGAAGTAGGTACCATCGGTGGTGGTACTGTTCTAGAACCACAAGGTGCCATGT<br>TGGACTTATTAGGTGTAAGAGGCCCACATGCTACCGCTCCTGGTACCAACGCACGTCAATTAGCAAGAATAGTTGCCTGTGCC<br>GTCTTGGCAGGTGAATTATCCTTATGTGCTGCCCTAGCAGCCGGCCATTTGGTTCAAAGTCATATGACCCACAACAGGAAACCT<br>GCTGAACCAACAAAACCTAACAATTTGGACGCCACTGATATAAATCGTTTGAAAGATGGGTCCGTCACCTGCATTAAATCCTGA | BY4741 genome |

**Supplementary Table 2.** Plasmids used for strain construction. Promoters are represented by "pr", terminators are represented by "t".

| Plasmid | Backbone | Type | Insert | Source |
|---|---|---|---|---|
| pDB96 | pDZ306 | Integrative plasmid (single) | GLT1-5xELbs-CYC180pr-24xPP7SL | [8] |
| pDB99 | pFA6-his3MX6 | Integrative plasmid (single) | pr2xBS-CYC180pr-Kozak-mKate2-tADH1-HIS3MX | [6] |
| pYTKmk47 | pYTK096 | Integrative plasmid (single) | prRPL18B-Msn2AD-EL222(AQTrip)-tENO2 | this work |
| pYTKmk48 | pYTK096 | Integrative plasmid (single) | prRPL18B-Msn2AD-EL222(WT)-tENO2 | this work |
| pYTKmk165 | pYTK096 | Integrative plasmid (single) | prRPL18B-Msn2AD-EL222(A79Q)-tENO2 | this work |
| pYTKmk167 | pYTK096 | Integrative plasmid (single) | prRPL18B-Mig1RD-EL222(A79Q)-tENO2 | this work |
| pYTKmk54 | pYTK096 | Integrative plasmid (multigene) | prRPL18B-Msn2AD-EL222(AQT)-tENO2;prTDH3-Mig1RD-EL222(wt)-tPGK1 | this work |
| pYTKmk90 | pYTK096 | Integrative plasmid (multigene) | prRPL18B-Msn2AD-EL222(AQT)-tENO2;prTDH3-Mig1RD-EL222(A79Q)-tPGK1 | this work |
| pYTKmk92 | pYTK096 | Integrative plasmid (multigene) | prRPL18B-Msn2AD-EL222(AQT)-tENO2;prHHF1-Mig1RD-EL222(A79Q)-tPGK1 | this work |
| pSO40 | | Integrative plasmid (backbone) | Backbone: *LEU2* marker gene, *LEU2* homology arms | this work |
| pSO41 | pSO40 | Integrative plasmid (multigene) | prRPL18B-VP16AD-CIB1-tPGK1--prRPL18B-LexA-CRY2PHR(W349R)-tENO2--prLexACYCmin-Venus-tADH1 | this work |
| pYTKmk110 | | Integrative plasmid (backbone) | Backbone: Hygromycin B resistance, HO-locus homology arms | this work |
| pYTKmk116 | pYTKmk110 | Integrative plasmid (multigene) | prRPL18B-dCas9-Mxi1-tENO2--pr5xBS-CYC180pr-LexABSsgRNA-tENO2 | this work |
| pYTKmk156 | pYTKmk110 | Integrative plasmid (multigene) | pr5xBS-CYC180-GAL7sgRNA-tENO2--prLexaCYCmin-dCas9-VPR-tENO1--prGAL7-mTurqoise-tPGK1 | this work |
| pYTKmk202 | pYTKmk110 | Integrative plasmid (multigene) | pr5xBS-CYC180-ycrtI-tENO1--prLexaCYCmin-ycrtE-tENO2--prLexaCYCmin-ycrtYB-tPGK1--prPGK1-HMG(t)-tTDH1 | this work |

**Supplementary Table 3.** Strains used in this study. Promoters are represented by “pr”, terminators are represented by “t”.

| Name | Genotype | Source | Data shown in Figure |
|---|---|---|---|
| BY4741 | MATa his3Δ1 leu2Δ0 met15Δ0 ura3Δ0 | Euroscarf | - |
| BY4742 | MATalpha his3Δ1 leu2Δ0 lys2Δ0 ura3Δ0 | Euroscarf | - |
| DBY73 | BY4741, his3Δ::pr5xBS-CYC180-Kozak-mKate2-tADH1t-HIS3MX(pDB60) | Nat. Comm | |
| DBY91 | BY4742, URA3::prMET25-tdPCP-NLS-tdmRuby3-tCYC1(pDB97) | Mol Cell. | - |
| SOY9 | BY4741, prGLT1Δ::HIS3-pr5xELbs-CYC180-24xPP7SL(pDB96) | this work | - |
| DBY175 | DBY73, URA3::prRPL18B-Msn2AD-EL222(AQTrip)-tENO2(pYTKmk47) | this work | Fig. 2c,d |
| DBY176 | DBY73, URA3::prRPL18B-Msn2AD-EL222(WT)-tENO2(pYTKmk48) | this work | Fig. 2c,d |
| DBY177 | DBY73, URA3::prRPL18B-Msn2AD-EL222(A79Q)-tENO2(pYTKmk165) | this work | Fig. 2c,d |
| DBY193 | BY4741, his3Δ::pr2xBS-CYC180pr-Kozak-mKate2-tADH1-HIS3MX(pDB99) | this work | - |
| DBY194 | DBY193, URA3::prRPL18B-Mig1AD-EL222(A79Q)-tENO2(pYTKmk167) | this work | Fig. 2f |
| DBY189 | SOY9, URA3::prRPL18B-Msn2AD-EL222(AQT)-tENO2;prTDH3-Mig1RD-EL222(wt)-tPGK1(pYTKmk54) | this work | - |
| DBY190 | DBY91/DB190 | this work | Fig. 3d,e |
| DBY170 | DBY73, URA3::prRPL18B-Msn2AD-EL222(AQT)-tENO2;prTDH3-Mig1RD-EL222(A79Q)-tPGK1(pYTKmk90) | this work | Fig. 4b,c |
| DBY172 | DBY73, URA3::prRPL18B-Msn2AD-EL222(AQT)-tENO2;prHHF1-Mig1RD-EL222(A79Q)-tPGK1(pYTKmk92) | this work | Fig. 4b,c |
| DBY179 | DBY73, URA3::prRPL18B-Msn2AD-EL222(AQT)-tENO2;prTDH3-Mig1RD-EL222(wt)-tPGK1(pYTKmk54) | this work | Fig. 4b,c |
| DBY183 | DBY170, LEU2::prRPL18B-VP16AD-CIB1-tPGK1--prRPL18B-LexA-CRY2PHR(W349R)-tENO2--prLexACYCmin-Venus-tADH1(pSO41) | this work | Fig. 5b-d |
| DBY185 | DBY183, HO::prRPL18B-dCas9-Mxi1-tENO2--pr5xBS-CYC180pr-LexABSsgRNA-tENO2(YTKmk116) | this work | Fig. 6b |
| DBY188 | DBY183, HO::pr5xBS-CYC180-GAL7sgRNA-tENO2--prLexaCYCmin-dCas9-VPR-tENO1--prGAL7-mTurqoise-tPGK1(pYTKmk156) | this work | Fig. 6d |
| DBY195 | DBY183, HO::pr5xBS-CYC180-ycrtI-tENO1--prLexaCYCmin-ycrtE-tENO2--prLexaCYCmin-ycrtYB-tPGK1--prPGK1-HMG(t)-tTDH1(YTKmk202) | this work | Fig. 7c,d |

**Supplementary Table 4.** Parameters used for model simulations.

| Parameter | Description | Activation (Fig. 2c,d) | Repression (Fig. 2e) |
|---|---|---|---|
| TFtot (molecules) | total cellular TF | 2000 / 1500 (IFFL) | 2000 / 20000 (IFFL) |
| kon (min-1 * (uW cm-2)-1) | light dependant VP-EL222 activation rate | 0.00606 (wt) / 0.0039878 (A79Q) / 0.15385 (AQT) | 0.0039878 |
| koff (min-1) | VP-EL222 dark-state reversion rate | 0.343 (wt) / 0.11324 (A79Q) / 0.02491 (AQT) | 0.11324 |
| kbasal (mRNA * min-1) | basal transcription rate | 0.012389 | 0.185 |
| kmax (mRNA * min-1) | maximal induced transcription rate | 30.323 | 0.184 |
| Kd (molecules) | TFon level required for achieving kmax / 2 | 1267 | 1032 |
| n (-) | hill coefficient | 3.268 | 3.268 |
| kdegR (min-1) | mRNA degradation rate | 0.042 | 0.042 |
| ktrans (proteins * min-1 * mRNA-1) | translation rate | 0.37 | 0.37 |
| kdegP (min-1) | protein degradation rate | 0.007 | 0.007 |
